# Selective Translation Orchestrates Key Signaling Pathways in Primed Pluripotency

**DOI:** 10.1101/2024.02.09.579580

**Authors:** Chikako Okubo, Michiko Nakamura, Masae Sato, Yuichi Shichino, Mari Mito, Yasuhiro Takashima, Shintaro Iwasaki, Kazutoshi Takahashi

## Abstract

Pluripotent stem cell (PSC) identities, such as differentiation and infinite proliferation, have long been understood within the frameworks of transcription factor networks, epigenomes, and signal transduction, yet remain unclear and fragmented. Directing attention toward translational regulation, as a bridge between these events, promises to yield new insights into previously unexplained mechanisms. Our functional CRISPR interference screening-based approach revealed that EIF3D maintains primed pluripotency through selective translational regulation. The loss of EIF3D disrupts the balance of pluripotency-associated signaling pathways, impairing primed pluripotency. Moreover, we discovered that EIF3D ensures robust proliferation by controlling the translation of various p53 regulators, which maintain low p53 activity in the undifferentiated state. In this way, selective translation by EIF3D tunes the homeostasis of the primed pluripotency networks, ensuring the maintenance of an undifferentiated state with high proliferative potential. Therefore, this study establishes a paradigm for selective translational regulation as a defining feature of primed PSC identity.

## Introduction

Pluripotent stem cells (PSCs) have the capacity for self-renewal under appropriate conditions while maintaining their distinct attributes, including differentiation potential and unlimited proliferation^1–5^. Pluripotency is categorized into two types: naïve, resembling the pre- implantation inner cell mass, and primed, akin to the post-implantation epiblast^6^. These categories differ in their specific requirements for self-renewal, differentiation capacity, and epigenetic status. Previous studies have shown that inhibiting multiple kinases helps sustain naïve pluripotency in both rodents and humans, suggesting a conserved, kinase-independent strategy across species^7–10^.

Conversely, shifting from kinase inhibition to specific growth factor stimuli enables naïve PSCs to transition into primed pluripotency, a state poised for differentiation into various somatic lineages^7,11^. Unlike the naïve state, the fate of primed pluripotency depends on a range of signaling inputs, including FGF, IGF, and TGFβ^12–14^. Thus, kinase signaling dynamics are crucial for the transition between these states and their ongoing maintenance. Paradoxically, the same growth factors that support primed pluripotency also initiate lineage-specific differentiation programs^15,16^. Maintaining a delicate balance between strong and weak kinase signaling is key to preserving the equilibrium between self-renewal and differentiation induction^17,18^. Although primed pluripotency, maintained by finely tuned signaling, is a significant research area in stem cell and developmental biology, the complex mechanisms governing this balance remain elusive.

The translation process, converting RNA into proteins, emerges as a critical element in cellular homeostasis and the study of primed pluripotency. While overall translation remains low during stem cell maintenance, differentiation cues actively enhance protein synthesis^19^. This shift in translation dynamics highlights the importance of translational control in dictating pluripotency and differentiation. Analysis of genes showing discrepancies between mRNA and protein levels has unveiled the critical role of context-dependent post-transcriptional regulation in maintaining primed pluripotency^20^. This indicates that translational modulation significantly influences primed pluripotent states, independent of transcriptional regulation. While several translational factors associated with primed pluripotency have been identified^21–23^, a comprehensive understanding of the role of translational regulation in stem cell homeostasis and fate determination remains elusive.

## Results

### EIF3D Essential for Human Primed Pluripotency

To investigate the complex mechanisms of primed pluripotency, we conducted genome-wide functional screening using the CRISPR interference (CRISPRi) platform^24,25^. This approach identified 1,686 genes that positively influence the self-renewal of primed human PSCs (Fig. 1a and Supplementary Table 1). Gene ontology (GO) analysis of these genes, critical for maintaining primed PSC identity, highlighted translation-related terms as significantly enriched (Fig. 1b).

**Fig. 1:**
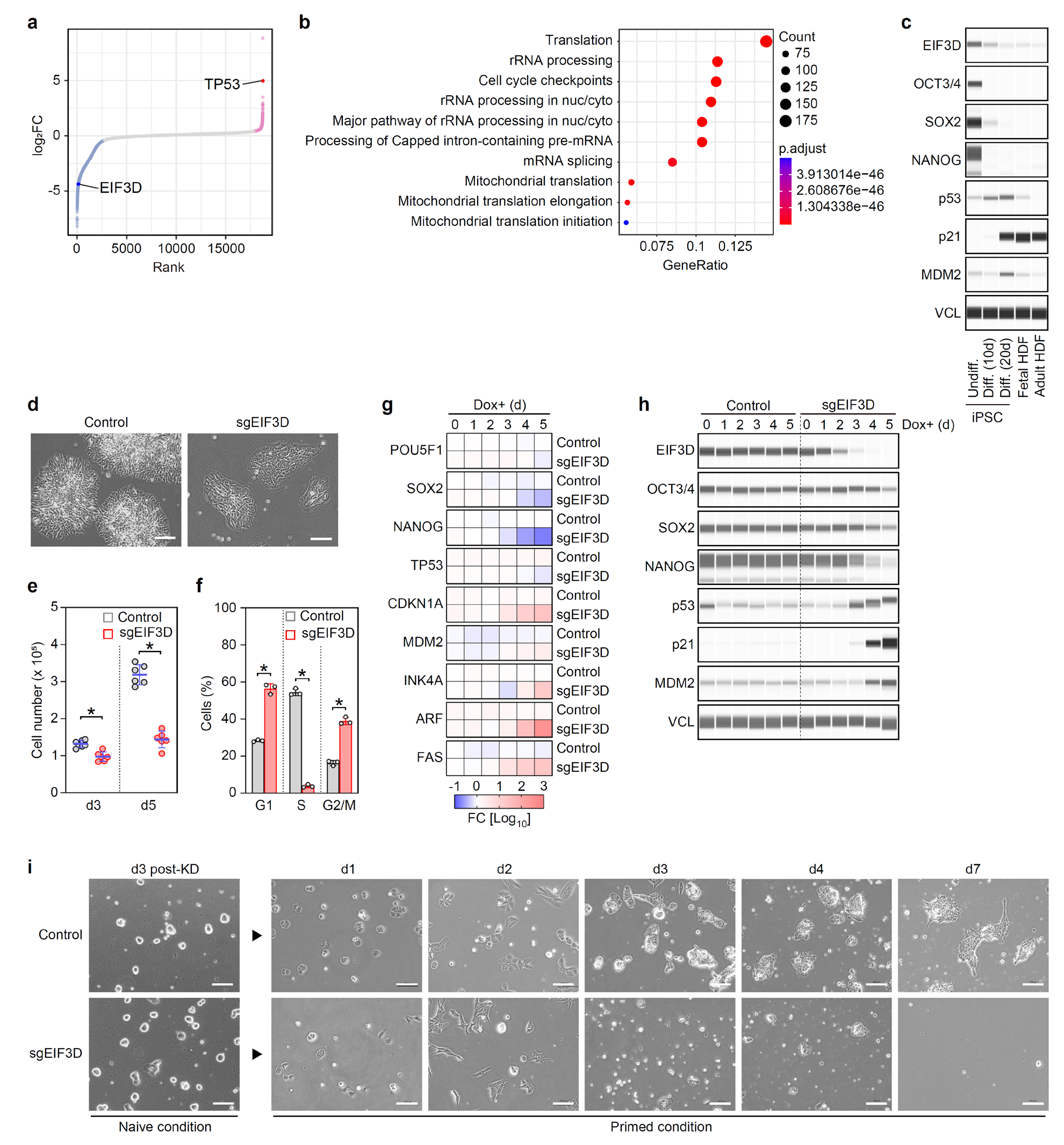
EIF3D is essential for maintaining primed pluripotency. **a**, Rank plot from CRISPRi screening. Red and blue dots represent genes significantly increased or decreased (±1 standard deviation (SD)) following 16 days of knockdown, respectively. n=3. See full list in Supplementary Table 1. **b**, Top gene ontology terms among 1,686 genes crucial for primed pluripotency maintenance. **c**, Protein expression in undifferentiated and differentiating PSCs (10 and 20 days post-FGF withdrawal) and HDFs. **d**, Representative images of Control and EIF3D KD iPSCs, 5 days post-KD induction. Scale bars: 100 μm. **e**, Cell counts on days 3 (p=4.70e-4) and 5 (p=3.53e-7) post-KD induction (mean ± SD, n=6). p-values determined via unpaired t-test. **f**, Cell cycle phase distribution (mean ± SD, n=3). G1: p=2.14e-3; S: p=4.43e-7; G2/M: p=1.99e-4, calculated by unpaired t-test. **g**, RNA expression of pluripotency and TP53- related genes during EIF3D KD. n=3. **h**, Expression of pluripotency and p53-associated proteins during EIF3D KD. **i**, Differentiation of naive PSCs to primed state. Induction of differentiation from naive to primed PSCs, 3 days post-Dox addition, by altering culture conditions. Representative images of specified cell lines and days are shown. Scale bars: 100 μm. See also Supplementary Fig. 1, 2.

We focused on one of the top-ranked translation regulators from the screening, the eukaryotic translation initiation factor subunit D (EIF3D), a cap-binding protein^26^. EIF3D is abundantly expressed in undifferentiated induced PSCs (iPSCs) compared to dermal fibroblasts (HDFs), and its levels sharply decrease upon differentiation (Fig. 1c). This expression pattern is similar to pluripotency factors like OCT3/4, SOX2, and NANOG, suggesting EIF3D’s potential role in primed PSCs.

To explore its function, we created doxycycline (Dox)-inducible EIF3D knockdown (KD) iPSC lines (Supplementary Fig. 1a). Following EIF3D KD, we observed morphological changes characterized by flattened colonies with indistinct edges and notably enlarged nuclei (Fig. 1d and Supplementary Fig. 1b, 1c). Additionally, EIF3D KD resulted in a marked reduction in cell numbers between days three and five post-induction (Fig. 1e). Investigating this phenotype, we measured DNA synthesis via 5-ethynyl-2’-deoxyuridine (EdU) incorporation, revealing a significant decrease in the S phase cell population and a corresponding increase in cells in the G1 and G2/M phases following EIF3D KD (Fig. 1f and Supplementary Fig. 1d-f). These results collectively suggest that EIF3D KD imposes growth arrest on primed human PSCs.

Moreover, EIF3D KD reduced the expression of transcripts encoding core pluripotency transcription factors, which are indicative markers of PSCs (Fig. 1g). The decrease in NANOG expression was more rapid and pronounced than that of POU5F1 (encoding OCT3/4) and SOX2. Protein expression analyses exhibited a similar trend (Fig. 1h and Supplementary Fig. 1g). Notably, transcriptional targets of p53, such as CDKN1A and MDM2 mRNAs, and their translation products, were significantly upregulated in EIF3D KD cells, akin to the changes seen in differentiated iPSCs induced by FGF withdrawal (Fig. 1c, g, h). Other hallmarks of senescence (INK4A and ARF) and cell death (FAS) markers were also elevated (Fig. 1g). Similar to iPSC differentiation, the absence of EIF3D led to increased p53 protein expression, while TP53 mRNA showed only modest changes (Fig. 1c, g, h, and Supplementary Fig. 1g). Overall, the KD phenotypes suggested that EIF3D loss diminishes the primed PSC identity.

Given these results, we explored the potential roles of increased p53 levels and reduced NANOG expression in the phenotypes arising from EIF3D KD. Introducing exogenous NANOG did not restore pluripotency marker expression or resolve the proliferative impairment (Supplementary Fig. 2a-d). Additionally, the concurrent KD of TP53 with EIF3D did not fully restore pluripotency marker expression (Supplementary Fig. 2e-h). However, these experiments showed partial recovery of cellular proliferation and a decrease in elevated p21 expression (Supplementary Fig. 2g, h). These findings suggest that analyzing hallmark genes associated with undifferentiated or differentiated states alone is insufficient to fully understand EIF3D’s role in pluripotency. Nonetheless, the data indicate the involvement of the p53-p21 pathway in regulating the proliferation of primed PSCs, although other pathways likely contribute to EIF3D- mediated self-renewal. In summary, the collective findings highlight the multifaceted role of EIF3D in maintaining primed pluripotency through complex molecular interactions.

Next, we investigated the differentiation of naïve PSCs into a primed state to further assess EIF3D’s role in maintaining primed pluripotency. Prior to transitioning to the primed state, we induced EIF3D KD in naïve PSCs for 3 days, which did not result in any observable abnormalities (Fig. 1i). However, when we altered the culture conditions from the kinase-inhibiting naïve state to the growth factor-rich primed state, the EIF3D KD naïve PSCs demonstrated an inability to differentiate into the primed state. This was marked by significant cell death within 4 days (Fig. 1i). Given this evidence of the inability to self-renew primed PSCs following EIF3D KD, we conclude that EIF3D is essential for maintaining primed pluripotency.

### Loss of EIF3D Diminishes Primed Pluripotency with Limited Impact on Three Germ Layer Specifications

We then conducted a comprehensive genome-wide transcriptome analysis to further investigate the underlying mechanisms from a broader perspective. To elucidate the cell fate changes in primed PSCs induced by EIF3D KD, we compared the global gene expression profiles of EIF3D KD cells with those of iPSCs differentiated through suppression of core transcription factors (Supplementary Fig. 3a-c), and iPSC-derived cells directed towards endoderm (EN), mesoderm (ME), and neuroectoderm (NE) lineages (Supplementary Fig. 3d).

Our findings revealed that EIF3D KD resulted in an incremental increase in differentially expressed genes (DEGs) over successive days compared to the controls (Fig. 2a and Supplementary Fig. 3e). Notably, despite significant changes in gene expression, the EIF3D KD profile showed less similarity to the corresponding comparatives (Fig. 2a and Supplementary Fig. 3f). Instead, it seemed to enter an independent state, marked by a lack of clear lineage commitment to any of the three germ layers. This was accompanied by the downregulation of key pluripotency and primed PSC markers, including ZIC2, CD24, and SFRP2 (Fig. 2a, b, and Supplementary Table 2) ^27–32^. GO analysis revealed that EIF3D KD upregulated genes associated with inconsistent differentiation terms, while genes related to cell cycle and division were downregulated (Supplementary Fig. 3g). These results align well with the observed EIF3D KD phenotypes, including growth retardation and loss of pluripotency, with minimal contribution to specific lineages.

**Fig. 2:**
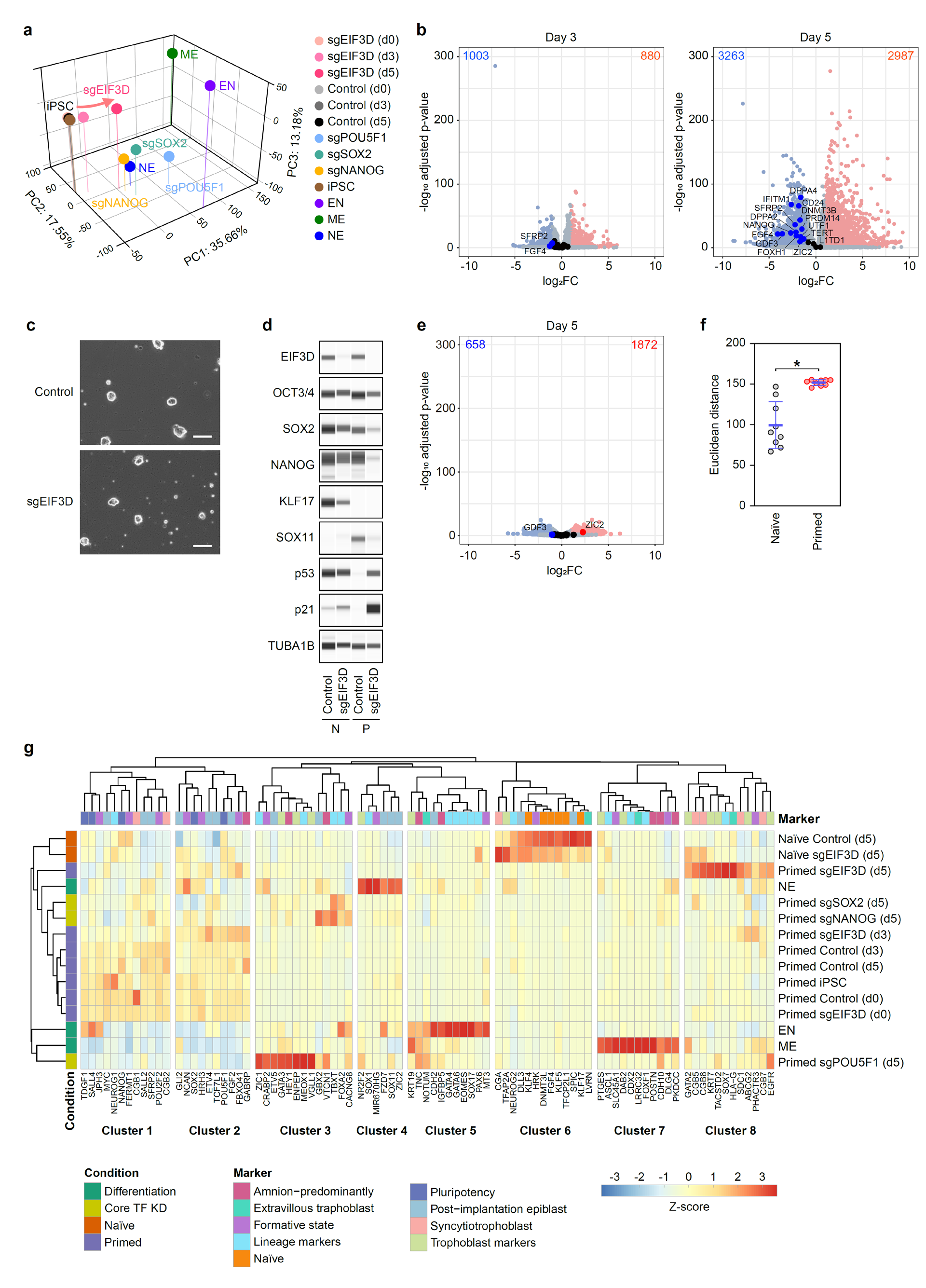
Transcriptome dynamics in primed PSCs following EIF3D loss. **a**, Principal component analysis (PCA) of RNA-seq data. Each dot represents the average value of replicates. n=3. **b**, Volcano plots displaying differentially expressed genes (DEGs) between control and EIF3D KD iPSCs, 3 and 5 days after Dox addition. Understated-colored dots represent DEGs. Highlighted-colored dots denote key pluripotency markers. Red and blue dots indicate genes significantly upregulated and downregulated, respectively (|log2FC| > 1, adjusted p < 0.05). n=3. See full gene list and FC in Supplementary Table 2. **c**, Representative images of control and sgEIF3D naïve PSCs, 5 days post-Dox addition. Scale bars: 100 μm. **d**, Protein expression in control and EIF3D KD naïve (N) and primed (P) PSCs, 5 days post-Dox addition. **e**, Volcano plot illustrating DEGs (|log2FC| > 1, adjusted p-value < 0.05) between control and sgEIF3D naïve PSCs, 5 days post-Dox addition. **f**, Euclidean distance between control and sgEIF3D in both naïve and primed PSCs (mean ± SD, n=9) (unpaired comparison of three each from control and sgEIF3D groups). p=5.84e-4, determined by unpaired t-test. **g**, Sample clustering from Fig. 2A in addition to control and sgEIF3D naïve PSCs, with emphasis on selected marker genes. n=3. See also Supplementary Fig. 3.

To further investigate the effects of EIF3D KD in primed PSCs, beyond the typical three germ layers derived from these cells, our study expanded to include marker genes related to naïve PSCs and the trophoblast, an earlier diverging fate^33^. EIF3D KD also appears to weaken naïve pluripotent signatures, marked by decreased expression of naïve PSC markers, though without noticeable morphological changes in naïve PSC colonies (Supplementary Fig. 2c, d). However, the impact of EIF3D KD on gene expression changes in naïve PSCs was less pronounced compared to those in the primed state (Fig. 2e, f). These findings underscore the significant role of EIF3D in the regulation of primed PSCs.

Cluster analysis of gene sets showed that EIF3D downregulation in primed PSCs resulted in a significant divergence from the typical primed PSC cluster, as evidenced by reduced expression of genes linked to pluripotency, post-implantation epiblast, and the formative state, an intermediary between naïve and primed pluripotency^34^, predominantly in clusters 1 and 2 (Fig. 2g).

A notable characteristic of EIF3D KD primed PSCs is the increased expression of trophoblast genes, primarily found in cluster 8 (Fig. 2g). This cluster is distinct from clusters 4, 5, and 7, which contain cells that have differentiated into the three germ layers. This suggests that EIF3D KD phenotypes disrupt primed pluripotency, leading to mismanagement of cell fate, such as trophoblast gene activation, rather than inducing straightforward differentiation in line with developmental logic. The evidence collectively indicates that EIF3D maintains primed pluripotency through mechanisms distinct from those of core transcription factors or the inhibition of specific lineage commitments.

### EIF3D Orchestrates Translation of Key Signaling Pathways in Primed Pluripotency

Based on global transcriptome data, we observed a lesser degree of change induced by EIF3D KD on day 3 compared to day 5 (Fig. 2a, b, and Supplementary Fig. 3e, f). To understand the initial response to EIF3D loss, we analyzed translation statuses on day 3 post-KD induction. Puromycin incorporation showed that EIF3D KD reduced de novo protein synthesis to 45% relative to control cells (Fig. 3a). Polysome profile analysis indicated a significant accumulation of the 80S ribosomal subunit with EIF3D KD, suggesting decreased translation initiation (Fig. 3b, c).

**Fig. 3:**
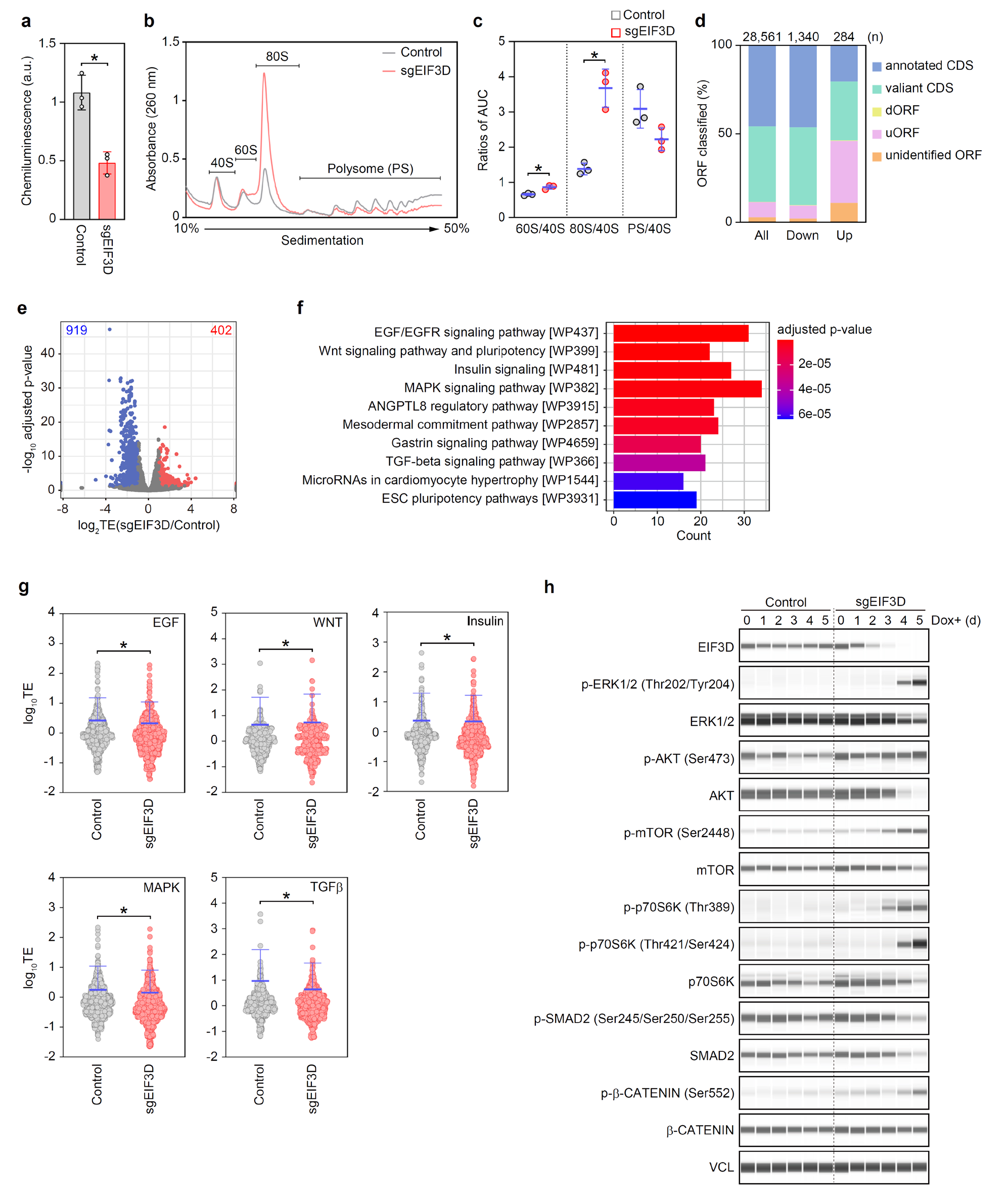
EIF3D-mediated selective translation regulates multiple signaling pathways. **a**, Quantification of de novo protein synthesis by detecting incorporated puromycin. n=3. p=6.75e-3 determined by unpaired t-test. **b**, Representative polysome profiles of sgEIF3D primed PSCs compared to control lines on day 3 post-Dox addition. **c**, Area under curve quantification for specified ribosomal fractions (mean ± SD, n=3). Ratios 60S/40S: p=0.011; 80S/40S: p=0.012; polysomes (PS)/40S: p=0.094, calculated using unpaired t-test. **d**, Categorization of ORFs with varying translation efficiency during EIF3D KD. Displayed are all ORFs translated in human iPSCs (All), and those downregulated (Down) or upregulated (Up) by EIF3D KD over 3 days. **e**, Volcano plot showing upregulated Differentially Translated Expressed Genes (uDTEGs, red) and downregulated DTEGs (dDTEGs, blue) (|log2FC| > 1, adjusted p < 0.05). See full gene list, FC, and adjusted p-value in Supplementary Table 3, 4. **f**, Pathway enrichment analysis of dDTEGs using WikiPathways. **g**, Beeswarm plots indicating log10TE of transcripts in specified pathways according to WikiPathways (WP437, WP399, WP481, WP382, and WP366 for EGF, WNT, Insulin, MAPK, and TGFβ pathways, respectively) (mean ± SD). EGF: p<0.0001; WNT: p=0.0262; Insulin: p<0.0001; MAPK: p<0.0001; TGFβ: p<0.0001, analyzed by Wilcoxon matched-pairs signed-rank test. See full gene list, FC, and adjusted p-value in Supplementary Table 5-9. **h**, Phosphorylation status of key proteins in the signaling pathways across the timeline of EIF3D KD.

To determine if EIF3D selectively regulates translation, we identified 28,561 open reading frames (ORFs) undergoing translation in human primed iPSCs, classified as follows: 45% as annotated coding sequences (CDS), 42% as variant CDS, 2% as unidentified ORFs, and 8% as upstream ORFs (uORFs) (see All in Fig. 3d). Our analysis revealed that EIF3D KD increased translation efficiency (TE) in 284 ORFs and decreased it in 1,340 ORFs. The increased TE group featured a higher percentage of uORFs (34.86%), whereas the reduced TE group showed no significant preference in ORF classification. These findings suggest that EIF3D is involved in regulating the translation of non-canonical ORFs in primed PSCs, though annotated ORFs like CDS comprise the majority of EIF3D targets.

Therefore, we subsequently examined the status of annotated ORF translation following EIF3D KD. Ribosome profiling demonstrated that EIF3D KD significantly altered the TE of 1,321 genes (increased in 402; decreased in 919), which we term differential translation efficiency genes (DTEGs). This suggests a selective translation regulation by EIF3D (Fig. 3e and Supplementary Table 3, 4). Pathway analysis indicated that downregulated DTEGs (dDTEGs) were linked to several signaling pathways, including EGF, WNT, insulin, MAPK, and TGFβ, all critical in maintaining pluripotency (Fig. 3f) ^35^. In contrast, upregulated DTEGs (uDTEGs) did not show significantly enriched terms. Given the EIF3D KD phenotype, which includes the loss of primed pluripotency, it is plausible that these pluripotency-related signaling pathways are implicated in dDTEG.

Following the pathway analysis results, we confirmed the TEs of EIF3D targets in each enriched pathway. The beeswarm plots revealed significant TE alterations in the transcripts of the EGF, WNT, Insulin, MAPK, and TGFβ pathways (Fig. 3g and Supplementary Table 5-9). Next, we assessed the phosphorylation statuses of key proteins in these signaling pathways, potentially regulated by EIF3D. EIF3D KD led to the hyperactivation of the MAPK, insulin (AKT, mTOR, and p70S6K), and WNT pathways, while simultaneously suppressing the TGFβ pathway through SMAD2 (Fig. 3h). These findings confirm that EIF3D KD disrupts the balance of multiple signaling activities in primed PSCs through selective translation regulation.

### EIF3D Inhibits p53 Protein Expression Through Selective Translation of p53 Regulators

Besides the dysregulation of multiple kinase pathways, there is a notable increase in p53 protein and subsequent activation of the p53 pathway due to EIF3D KD (Fig. 1h and Supplementary Fig. 1h, 2g). However, ribosome profiling data revealed no significant change in p53 translation efficiency following EIF3D KD (Likelihood ratio test, Fold Change=1.01, adjusted p=0.92). This led us to hypothesize about an indirect regulatory mechanism. To deepen our understanding, we conducted a comparative analysis between DTEGs and a compilation of post-transcriptional regulators of p53 protein expression^36^. Setting a TE threshold of 1.5-fold change, EIF3D KD resulted in significant translation dysregulation of 207 out of 818 genes, accounting for 25.3% of the list (Fig. 4a and Supplementary Table 10). We also confirmed the protein reduction of p53 regulators in EIF3D KD primed PSCs (Fig. 4b), suggesting that EIF3D indirectly influences p53 protein expression by regulating the translation of its regulators.

**Fig. 4:**
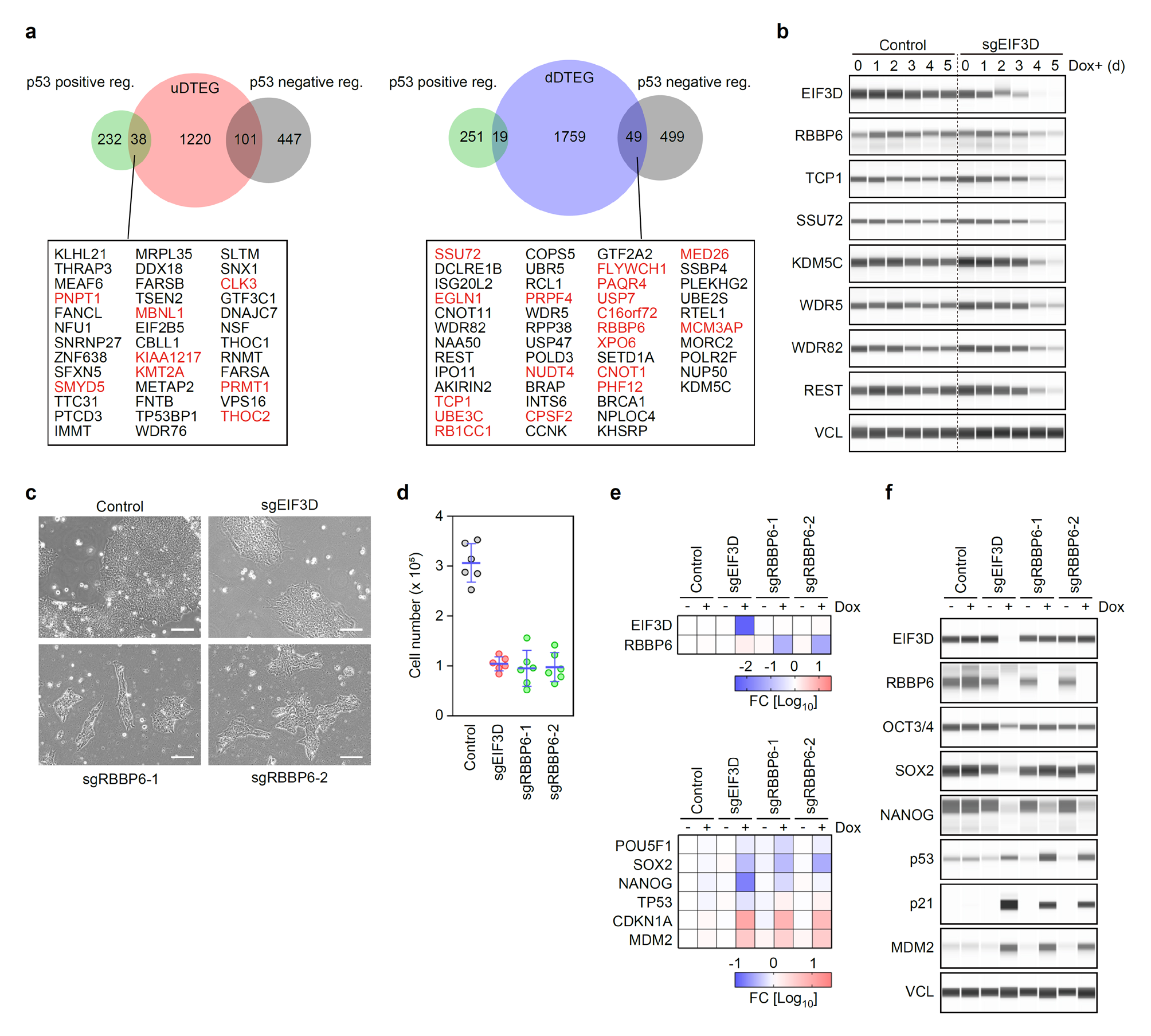
Indirect regulation of p53 protein through EIF3D-mediated selective translation. **a**, Venn diagrams display the overlap between DTEGs with significant Translation Efficiency (TE |log2FC| > 0.58) and p53 regulators. Genes with higher TE (|log2FC| > 1) are highlighted in red. See full gene list, FC, and adjusted p-value in Supplementary Table 10. **b**, Protein expression of p53 regulators translationally controlled by EIF3D. High TE (|log2FC| > 1) includes RBBP6, SSU72, and TCP1; moderate TE (|log2FC| > 0.58) includes KDM5C, WDR5, WDR82, and REST. **c**, Representative images of specified cells, 5 days post-KD induction. Scale bars: 100 μm. **d**, Cell counts of the cells depicted in Fig. 4c (mean ± SD, n=6). **e**, Relative gene expression in cells from Fig. 4c. Values normalized to GAPDH and compared to control without Dox. n=3. **f**, Protein expression of specified proteins in cells from Fig. 4c.

As an example, we identified RBBP6, a dDTEG, previously reported as a negative regulator of p53 protein stability^37^. As expected, RBBP6 KD mimicked EIF3D KD phenotypes, including cell growth defects with morphological changes, reduced expression of pluripotency markers, increased p53 proteins, and elevated expression of CDKN1A/p21 and MDM2 (Fig. 4c- f). These findings indicate that EIF3D plays a role in modulating p53 protein expression by controlling the translation of its regulators.

## Discussion

This study demonstrates that EIF3D is crucial in sustaining primed pluripotency through selective translation, which finely balances kinase signaling and suppresses the p53 pathway. EIF3D’s role, inclusive of selective translation, has been recognized in relation to oncogenic and stress responses. Recent research has clarified EIF3D’s influence on the translation regulation of the MAPK pathway in common cell lines, such as 293T and HeLa^38,39^. These prior findings bolster our current results, showing that EIF3D’s translation control can modulate specific signaling pathways’ activities.

Maintaining low p53 activity is essential in undifferentiated PSCs, though its exact regulatory mechanism was previously unclear^40^. Our study sheds light on EIF3D’s indirect inhibition of p53 by targeted translation modulation of p53 regulators. Moreover, the marked increase of p53 protein in EIF3D KD, combined with the inverse correlation between increased p53 protein and decreased EIF3D expression during PSC differentiation, strongly suggests EIF3D’s pivotal role in suppressing p53.

EIF3D KD led to a notable phenotype in primed PSCs, yet its effect was limited in the naïve state. This difference might be due to the abundant expression and EIF3D-insensitive regulation of p53 protein in naïve PSCs. Additionally, kinase pathway-independent self-renewal in kinase-inhibiting conditions might make naïve PSCs less susceptible to EIF3D’s translation regulation. Although further investigation is needed, the significant phenotype observed in EIF3D KD naïve PSCs exposed to growth factor-rich media for primed PSCs supports this theory.

Research on transcription factor networks and signaling pathways has greatly enhanced our understanding of pluripotency. This study highlights the significance of translation regulation as a link between these two aspects in pluripotency. Our CRISPRi screening indicates that, besides EIF3D, other translational regulators may play roles in maintaining primed pluripotency, suggesting that deeper exploration into translational control will offer more insights into pluripotency’s fundamental nature.

## Methods

### Cell Culture

Human iPSC lines (WTB6 and 1B4 are gifts from Bruce R. Conklin of the Gladstone Institutes) were maintained on tissue culture plates coated by iMatrix 511 silk (Matrixome) using StemFit AK02N media (Ajinomoto), as previously described^41^. For passaging, cells were washed once with Dulbecco’s Phosphate Buffered Saline (D-PBS, Nacalai Tesque) and incubated in TrypLE Express (Thermo Fisher Scientific) for 10 min at 37°C. Subsequently, cells were dissociated into single cells and washed in Dulbecco’s Modified Eagle Medium/Ham’s F-12 (DMEM/F-12, WAKO) containing 0.1% bovine serum albumin (BSA, WAKO). After cell counting and centrifugation, the cells were resuspended in StemFit AK02N media supplemented with 1.67 μg/mL iMatrix-511 silk and 10 μM Y-27632 (Nacalai Tesque). G-banding tests conducted by Nihon Gene Laboratories confirmed that all PSC lines used in this study showed no significant karyotypic abnormalities. Human dermal fibroblasts (HDFs) derived from fetal (HDF1419, Cell Applications) and adult (TIG- 120, a gift from Kazuhiko Kaji) donors were maintained in DMEM (Nacalai Tesque) with 10% fetal bovine serum (FBS, Cosmo Bio) and used within six passages for the study. Routine testing confirmed the absence of mycoplasma infection.

### Transposon-Mediated Gene Transfer

We transfected 1 μg of a plasmid containing the inverted terminal repeats of either PiggyBac (PB) or Sleeping Beauty (SB) transposons, together with 0.5 μg of a plasmid encoding a hyperactive PB transposase (hyPBase) or SB transposase (SB100X), into 5 × 10^5^ human PSCs. This was accomplished using the P3 Primary Cell 4D-Nucleofector X Kit S (Lonza) and Program CA-137 on the 4D Nucleofector device (Lonza). Two days post-transfection, the transfectants were selected with the appropriate drug until non-transfected cells were completely eradicated. Subsequently, single cell-derived colonies that uniformly expressed the transduced fluorescent protein were isolated and expanded.

### CRISPR Interference (CRISPRi)

To generate inducible CRISPRi iPSC lines targeting a specific gene, we introduced a vector containing U6 promoter-driven sgRNA along with CAG promoter-driven fluorescence protein and a drug resistance marker into the 1B4 human iPSC line (P23-28) using the PB-mediated gene transfer method previously described^24^. Starting on day 2 post-transfection, drug selection was initiated and continued until non-transfected cells were eliminated. Subsequently, single cell-derived colonies uniformly expressing the fluorescent protein were isolated and expanded. To induce knockdown, we administered 1 μg/mL of doxycycline (Dox, WAKO) for the specified duration. Knockdown clones within 20 passages post-subcloning were utilized for the study. The spacer sequences are listed in Supplementary Table 11.

### Endoderm Differentiation

Endoderm differentiation was conducted as previously described, with minor modifications^42,43^. 1B4 iPSCs (P35, 36, and 37) were seeded at a density of 1 × 10^6^ cells per well in iMatrix 511- coated 6-well plates using StemFit AK02N media, supplemented with 10 μM Y-27632. The following day, cells were washed once with DMEM/F-12 and the media was replaced with Differentiation Media 1 (DM1) consisted of DMEM/F-12 (Thermo Fisher Scientific), 2% B27 supplement (Thermo Fisher Scientific), 1% MEM Non-Essential Amino Acids (NEAA, Thermo Fisher Scientific), and 0.1 mM 2-mercaptoethanol (2-ME, Thermo Fisher Scientific), supplemented with 100 ng/mL Activin A (Nacalai Tesque), 3 μM CHIR99021 (Nacalai Tesque), 20 ng/mL bFGF (Nacalai Tesque), and 50 nM PI-103 (Cayman Chemical). After 24 hours, the cells were washed with DMEM/F-12 and the medium was replaced with DM1 supplemented with 100 ng/mL Activin A and 250 nM LDN193189 (Stemgent). Two days later, following another wash with DMEM/F-12, the cells were cultured in DM1 with 100 ng/mL Activin A for an additional 48 hours.

### Mesoderm Differentiation

Directed differentiation to mesoderm was carried out with minor modifications from previously described methods^43,44^. A day prior to differentiation, 1B4 iPSCs (P35, P36, and P37) were seeded at a density of 1 × 10^6^ cells per well in iMatrix 511-coated 6-well plates using StemFit AK02N media supplemented with 10 μM Y-27632. The following day, cells were washed once with DMEM/F-12 and then cultured in DM1 medium containing 30 ng/mL Activin A, 40 ng/mL BMP4 (Peprotech), 6 μM CHIR99021, 20 ng/mL bFGF, and 100 nM PIK-90 (MedChemExpress) for 24 hours. Subsequently, after a wash with DMEM/F-12, the medium was replaced with DM1 supplemented with 40 ng/mL BMP4, 1 μM A83-01, and 4 μM CHIR99021, and cells were maintained for an additional 48 hours. Then, the cells were washed once more with DMEM/F-12 and then cultured in DM1 medium supplemented with 40 ng/mL BMP4 for another 48 hours.

### Ectoderm Differentiation

Neuroectoderm differentiation was conducted as previously described^45,46^. A day prior to induction, 1B4 iPSCs (P35, 36, and 37) were seeded at a density of 1 × 10^6^ cells per well in iMatrix 511- coated 6-well plates, using StemFit AK02N medium supplemented with 10 μM Y-27632. The following day, the cells were washed once with DMEM/F-12 and then cultured in Glasgow’s MEM (WAKO) containing 8% Knockout Serum Replacement (KSR, Thermo Fisher Scientific), 1 mM sodium pyruvate (Sigma-Aldrich), 1% MEM Non-Essential Amino Acids (NEAA), 0.1 mM 2-ME, 1 μM A83-01, and 250 nM LDN193189. This was maintained for five days, with daily media changes.

Generation and Maintenance of Naïve PSCs Primed PSCs were converted to a naïve pluripotent state as previously described^11^. Prior to conversion, we maintained primed PSCs on γ-ray irradiated primary mouse embryonic fibroblasts (MEFs) in DFK20 media, composed of DMEM/F-12, 20% KSR, 1% NEAA, 0.1 mM 2-ME, and 4 ng/mL bFGF. For harvesting, cells were treated with CTK solution (ReproCELL) and dissociated into single cells. We then seeded 1.5 × 10^5^ primed PSCs onto inactivated MEFs in a well of a 6- well plate using DFK20 media supplemented with 10 μM Y-27632. The cells were incubated at 37°C in a hypoxic environment (5% O2). The following day, the media was replaced with NDiff227 (Takara) supplemented with 1 μM PD325901 (Stemgent), 10 ng/mL LIF (EMD Millipore), and 1 mM Valproic acid (WAKO). After three days, we switched the media to PXGL, consisting of NDiff227 supplemented with 1 μM PD325901, 2 μM XAV939 (WAKO), 2 μM Gö6983 (Sigma-Aldrich), and 10 ng/mL LIF. Upon the emergence of round-shaped colonies, the cells were dissociated using a 1:1 mixture of TrypLE Express and 0.5 mM EDTA, and then plated onto fresh inactivated MEF feeders in PXGL media containing 10 μM Y-27632. We replaced the media daily and passaged the cells every 3–5 days. The cells were utilized for assays after a minimum of 30 days post-conversion.

### Differentiation of Naïve PSCs to the Primed State

Prior to differentiating naïve PSCs into a primed state, we treated the cells, which were grown in PXGL media on iMatrix 511-coated plates, with Dox for 3 days. Subsequently, the media was replaced with StemFit AK02N, also supplemented with Dox, and the cells were incubated under normoxic conditions (20% O2). The cells were passaged every 4 days.

### RNA Isolation and Reverse-Transcription Polymerase Chain Reaction

Cells were washed once with D-PBS and lysed using QIAzol reagent (QIAGEN). Total RNA was extracted using the Direct-zol RNA Miniprep kit (Zymo Research), including on-column genomic DNA digestion as per the provided instructions. For reverse transcription (RT), one microgram of RNA was utilized, employing the ReverTra Ace qPCR RT Master Mix (TOYOBO). Quantitative RT-PCR was conducted with gene-specific primers (refer to Supplementary Table 11) using either THUNDERBIRD Next SYBR qPCR Mix (TOYOBO) or TaqMan assays (Thermo Fisher Scientific) with TaqMan Universal Master Mix II, no UNG (Thermo Fisher Scientific) on a QuantoStudio 5 Real-Time PCR System (Applied Biosystems). Raw Ct values were normalized against ACTB or GAPDH expression using the delta-delta Ct method. Relative expression was then calculated as fold-change relative to the control.

### Size-Based Protein Analysis

Cells were washed once with D-PBS and lysed using RIPA buffer (Sigma-Aldrich) supplemented with a protease inhibitor cocktail (Sigma-Aldrich). The crude lysates were centrifuged at 15,300 × g for 15 min at 4°C, and the cleared supernatant was transferred to a new tube. The concentration of the cleared lysate was measured using a Pierce BCA Protein Assay Kit (Thermo Fisher Scientific) and an EnVision 2104 plate reader (Perkin Elmer), following previously described methods. For quantitative and specific detection of target proteins, we utilized either a Wes or Jess automated capillary electrophoresis platform (ProteinSimple) with 12-230 kDa or 60-440 kDa Separation Modules (ProteinSimple). We loaded 2 μg of cell lysate per detection, along with the following antibodies: mouse monoclonal anti-OCT3/4 (1:500, Santa Cruz Biotechnology), goat polyclonal anti-SOX2 (1:40, R&D Systems), goat polyclonal anti-NANOG (1:40, R&D Systems), mouse monoclonal anti-p53 (DO-7) (1:200, Novus Biologicals), goat polyclonal anti-p53 (1:100, R&D Systems), rabbit monoclonal anti-p21 (1:50, Cell Signaling Technology), rabbit monoclonal anti-MDM2 (1:50, Cell Signaling Technology), rabbit polyclonal anti-EIF3D (1:250, Proteintech), rabbit polyclonal anti-SOX11 (1:500, Proteintech), rabbit polyclonal anti-KLF17 (1:200, Sigma-Aldrich), rabbit monoclonal anti-ERK1/2 (1:50, Cell Signaling Technology), rabbit monoclonal anti-phospho-ERK1/2 (Thr202/Tyr204) (1:50, Cell Signaling Technology), rabbit monoclonal anti- SMAD2 (1:50, Cell Signaling Technology), rabbit polyclonal anti-phospho-SMAD2 (Ser245/250/255) (1:50, Cell Signaling Technology), rabbit monoclonal anti-mTOR (1:50, Cell Signaling Technology), rabbit monoclonal anti-phospho-mTOR (Ser2448) (1:50, Cell Signaling Technology), rabbit monoclonal anti-AKT (1:50, Cell Signaling Technology), rabbit monoclonal anti-phospho-AKT (Ser473) (1:50, Cell Signaling Technology), rabbit polyclonal anti-p70S6K (1:50, Cell Signaling Technology), rabbit polyclonal anti-phospho-p70S6K (Thr389) (1:50, Cell Signaling Technology), rabbit polyclonal anti-phospho-p70S6K (Thr421/Ser424) (1:50, Cell Signaling Technology), rabbit monoclonal anti-β-Catenin (1:50, Cell Signaling Technology), rabbit monoclonal anti-phospho-β-Catenin (Ser552) (1:50, Cell Signaling Technology), mouse monoclonal anti-puromycin (1:20, Developmental Studies Hybridoma Bank), rabbit polyclonal anti-RBBP6 antibody (1:50, Sigma-Aldrich), rabbit polyclonal anti-TCP1 (1:200, Proteintech), rabbit polyclonal anti-SSU72 (1:100, Proteintech), rabbit polyclonal anti-REST (1:100, Proteintech), rabbit polyclonal anti-KDM5C (1:250, Proteintech), rabbit polyclonal anti-WDR5 (1:100, Proteintech), rabbit polyclonal anti-WDR82 (1:50, Proteintech), rabbit monoclonal anti-VINCULIN (1:250, Cell Signaling Technology), and rabbit polyclonal anti-alpha tubulin (1:200, Proteintech). Data visualization and analysis were conducted using Compass for SW6.0 software (ProteinSimple).

### Immunocytochemistry

The cells were washed once with D-PBS and fixed with 4% paraformaldehyde (Nacalai Tesque) for 15 min at room temperature. They were then blocked in D-PBS containing 1% BSA, 2% normal donkey serum (Sigma-Aldrich), and 0.2% Triton X-100 (Teknova) for 45 min at room temperature. Subsequently, the fixed cells were incubated overnight at 4°C with primary antibodies diluted in D-PBS containing 1% BSA. Following this, the cells were washed with D-PBS and incubated for 45 min at room temperature in 1% BSA containing fluorescence-conjugated secondary antibodies and 1 μg/mL Hoechst 33342 (Thermo Fisher Scientific). After a final wash in D-PBS, fluorescence was detected using a BZ-X810 imaging system (KEYENCE). Merged images were generated using a BZ-X Analyzer (KEYENCE). Nuclear size was quantified by analyzing Hoechst images with a Hybrid Cell Count Module (KEYENCE). The antibodies and their dilutions were as follows: mouse monoclonal anti-OCT3/4 (1:200), goat polyclonal anti-SOX2 (1:100), goat polyclonal anti-NANOG (1:100), goat polyclonal anti-p53 (1:200), rabbit monoclonal anti-p21 (1:400), Alexa 647 Plus anti-mouse IgG (1:500, Thermo Fisher Scientific), Alexa 647 Plus anti-rabbit IgG (1:500, Thermo Fisher Scientific), and Alexa 647 Plus anti-goat IgG (1:500, Thermo Fisher Scientific).

### Puromycin Incorporation

After washing the cells grown in three wells twice with pre-warmed D-PBS, we added StemFit AK02N media containing 100 μg/mL Cycloheximide (CHX, Sigma-Aldrich) to one well, and StemFit AK02N media alone to the other two wells. Following a 10-min incubation at 37°C, we added 1 μM puromycin to one well containing CHX-treated cells and to one of the two non-treated wells, then continued incubation for 30 min at 37°C. Post-incubation, cells were washed with ice-cold D-PBS and lysed using RIPA buffer, supplemented with a protease inhibitor cocktail. Subsequently, the samples underwent Size-based protein analysis as described previously.

### Cell-Cycle Analysis

As previously described^47^, we conducted cell-cycle analysis using the Click-iT EdU Alexa Fluor 647 Flow Cytometry Assay Kit (Thermo Fisher Scientific). Cells seeded at a density of 5 × 10^5^ cells per well in a 6-well plate were cultured for five days in StemFit AK02N media supplemented with Dox. Subsequently, the cells were incubated in media containing 10 µM 5-ethynyl-2′- deoxyuridine (EdU) for 135 min at 37°C. The cells were then harvested and washed with 1% BSA. Following centrifugation, the cells were resuspended in Click-iT fixative and incubated for 15 min at room temperature. After washing the fixed cells with 1% BSA, they were permeabilized with 1× Click-iT Perm/Wash reagent for 15 min at room temperature. For EdU detection, we added D- PBS containing Copper Protectant, Alexa Fluor 647 picolyl azide, and 1× Click-iT EdU buffer additive to the cell suspension. The samples were then washed with 1× Click-iT Perm/Wash reagent and stained with 1 μg/mL FxCycle Violet (Thermo Fisher Scientific) for 30 min at room temperature. We analyzed 1 × 10^4^ cells using a FACS Aria II (BD Biosciences) and BD FACSDiva software (BD Biosciences). EdU (detected with Alexa 647) and DNA (detected with FxCycle Violet) were analyzed using APC (650/660 nm) and Pacific Blue (405/455 nm) filters, respectively. Data analysis was conducted using FlowJo software (FlowJo LLC).

### Genome-wide CRISPRi Screens

Ten micrograms of the genome-wide CRISPRi library hCRISPRi-v2 (courtesy of Jonathan Weissman: Addgene, #83969) along with 3.75 μg of psPAX2 (courtesy of Didier Trono: Addgene, #12260) and 1.25 μg of pMD2.G (courtesy of Didier Trono: Addgene, #12259) were transfected into 293T/17 cells (P27, ATCC) using TransIT-Lenti Transfection Reagent (Mirus). Cells, plated at 5 million per 100-mm collagen I-coated dish, were transfected the day before^25^. Two days post-transfection, the virus-containing supernatant was filtered through a 0.45-μm pore size PVDF filter (Millipore), and lentiviral particles were concentrated using the Lenti-X Concentrator (Takara) as per the instructions. The lentivirus was then infected into 1B4 iPSCs (P23) at a multiplicity of infection (MOI) of <0.4 (as determined by TagBFP fluorescence in the lentiviral vector) to achieve coverage of >1,000×. Three days post-infection, cells were selected with 1.5 μg/mL puromycin until all non-infected cells perished. Subsequently, the cells were plated at 10 million per 150-mm dish in StemFiT AK02N containing 10 μM Y-27632 and iMatrix-511 silk. The following day, the media was replaced with StemFiT AK02N supplemented with Dox. Cells were split every two to three days, maintaining a minimum of 100 million cells, corresponding to a 1,000× coverage. Cells maintained without Dox (day 0) and those with Dox for 16 days were harvested, and SSEA-5 (+) cells were collected using an autoMACS Pro Separator (Miltenyi biotec). Genomic DNA was purified from at least 100 million cells of each sample using NucleoSpin Blood XL (Takara) or QIAamp DNA Blood Midi Kit (QIAGEN). The purified DNA was digested overnight with SbfI-HF (New England Biolabs) and separated on a 0.8% TAE agarose gel. Post-electrophoresis, DNA fragments ranging from 350 to 700 bp were excised from the gel and purified using a QIAGEN gel extraction kit (QIAGEN). PCR and library preparation were conducted as previously described^25^. The libraries were sequenced using a NextSeq 500/550 High Output v2 Kit (Illumina) with custom primers, following the manufacturer’s protocol. Reads were aligned to the hCRISPRi- v2 sequences, counted, and analyzed using MAGeCK (version 0.5.9.5), then visualized using the MAGeCK flute (version 1.12.0) package in R (version 4.1.1) ^48,49^. GO analysis was conducted using clusterProfiler (version 4.2.2) ^50,51^.

### Polysome Profiling

The method used for polysome fractionation was based on a previously described method with minor modifications^52^. A single semiconfluent well of a 10-cm dish containing either control or sgEIF3D iPSCs was placed on a CoolBox XT Workstation (Biocision) to maintain a temperature of 4°C. This was followed by one gentle wash with 5 mL of ice-cold DPBS. The cells were then gently scraped and dissociated in 0.6 mL of ice-cold lysis buffer, consisting of 20 mM Tris-HCl (pH 7.5), 150 mM NaCl, 5 mM MgCl2 (Nacalai Tesque), 1 mM dithiothreitol (DTT, WAKO), a protease inhibitor cocktail, 100 μg/mL cycloheximide (CHX), 100 μg/mL chloramphenicol, and 1% Triton X-100. The cell suspension was collected into a pre-chilled 1.5-mL DNA LoBind Tube (Eppendorf). The lysate was incubated for 15 min on ice with 25 units/mL Turbo DNase (Thermo Fisher Scientific) before centrifugation at 20,000 × *g* for 10 min at 4°C. The cleared supernatant was then transferred to a fresh 1.5-mL tube. The samples were rapidly frozen using liquid nitrogen and stored at -80°C.

A continuous sucrose gradient ranging from 10% to 45% was prepared using 10% and 45% sucrose solutions (Sigma-Aldrich) in a 14 × 95 mm polyclear tube (Seton). The gradient was created in the presence of 100 μg/mL CHX and 1 mM DTT in polysome buffer (25 mM Tris-HCl, pH 7.5, 150 mM NaCl, and 15 mM MgCl2) using the Biocomp Gradient Master program (Biocomp). Thawed cell lysates were measured for RNA concentration using the Qubit RNA BR Assay Kit (Thermo Fisher Scientific). A consistent volume of cell lysate containing 40 μg of RNA from each sample (300 μL) was layered onto the continuous sucrose gradient. The polysomes were separated by centrifugation in a himac ultracentrifuge using a P40ST rotor (himac) at 36,000 rpm for 2.5 hours at 4°C. The relative RNA abundance in ribosomal subunits, monosomes, and polysomes was detected using a 254-nm ultraviolet light with the Biocomp Piston Gradient Fractionator (Biocomp). AUC were calculated using GraphPad Prism 8.0.2.

### RNA Sequencing

Cells were lysed using QIAzol reagent, and total RNA was purified according to the protocol mentioned earlier. RNA quality was assessed using an Agilent RNA6000 Pico Kit on a Bioanalyzer 2100 (Agilent). The library preparation and subsequent analysis were carried out following methods outlined in previous studies^53,54^. Briefly, 100 ng of DNase-treated total RNA was used for library preparation with the Illumina Stranded Total RNA Prep Ligation with Ribo-Zero Plus kit (Illumina). The libraries were evaluated using an Agilent High-Sensitivity DNA Kit (Agilent) and then sequenced using either a NextSeq 500/550 High Output v2 Kit (Illumina), NextSeq 1000/2000 P2 Reagents (100 cycles) v3 (Illumina), or HiSeq X (Illumina). The adapter sequence was trimmed using cutadapt-1.12^55^. Reads mapping to ribosomal RNA were excluded using SAM tools (version 1.10) ^56^ and Bowtie 2 (version 2.2.5) ^57^. Reads were aligned to the hg38 human genome using STAR Aligner (version 2.7.10b) ^58^. Quality checks were performed using RSeQC (version 4.0.0) ^59^. Reads were counted with HTSeq (version 0.13.5) ^60^ using the GENCODE annotation file (version 35) ^61^. Counts were normalized using DESeq2 (version 1.34.0) in R (version 4.1) ^62^. The DESeq2 package was also used to perform Wald tests. PCA and heatmaps were generated using prcomp and pheatmap, respectively. GO analysis was conducted and visualized using clusterProfiler^50,51^.

### Ribosome Profiling

Ribosome profiling was conducted as previously outlined^63,64^. Cells were lysed in a buffer containing 20 mM Tris-HCl (pH 7.5), 150 mM NaCl, 5 mM MgCl2, 1 mM DTT, 1% Triton X-100, 100 μg/mL chloramphenicol, and 100 μg/mL CHX, followed by a 15-minute DNase treatment on ice. RNA concentrations in the lysate were measured using the Qubit RNA BR assay kit (Thermo Fisher Scientific). We treated 10 μg of RNA with RNase I (Epicentre) for 45 min at 25°C. The ribosome footprint RNA was then concentrated via ultracentrifugation using a sucrose cushion (20 mM Tris-HCl, pH 7.5, 150 mM NaCl, 5 mM MgCl2, 1 mM DTT, 20 U/mL SUPERase-In (Thermo Fisher Scientific), 1 M Sucrose, 100 μg/mL chloramphenicol, and 100 μg/mL CHX). The resulting pellets were resuspended in pellet buffer (20 mM Tris-HCl, pH 7.5, 300 mM NaCl, 5 mM MgCl2, 1 mM DTT, 1% Triton X-100, and 20 U/mL SUPERase-In) and purified using the Direct-zol RNA Microprep kit (Zymo Research). The RNA samples were separated by electrophoresis, and fragments ranging from 17-34 nt were excised and purified using Dr. GenTLE Precipitation Carrier (Takara). These purified ribosome footprint RNAs were ligated with linker oligonucleotides containing an inner index sequence and a unique molecular identifier (UMI), followed by rRNA depletion using riboPOOLs for Ribo-seq (siTOOLs). The residual RNAs were reverse transcribed using ProtoScript II (New England Biolabs) and circularized with circLigase2 (Epicentre). The cDNA templates were amplified using Phusion polymerase (New England Biolabs) with index- sequenced primers.

To calculate translational efficiency, corresponding RNA-seq experiments were performed using RNA extracted from the lysis buffer. We utilized TRIZOL LS reagent (Thermo Fisher Scientific) and the Direct-zol RNA Microprep kit for RNA extraction. The RNA-seq libraries were prepared as instructed by manufacturer’s protocol, except using RiboPOOLs for RNA-seq (siTOOLs) in the rRNA depletion step and xGen UDI-UMI Adapters (Integrated DNA Technologies) in adapter ligation step. The cDNA libraries were sequenced following the RNA sequencing protocol. The reads were demultiplexed using the inner index and adapters were removed using fastp (version 0.22.0) and fastx-split^65^. To filter out reads mapping to rRNA, Bowtie2 and SAMtools were used. The remaining reads were aligned to the human genome (hg38) using STAR (version 2.7.10b), and duplicates were removed based on UMI using bam- suppress-duplicates. Quality control statistics were calculated using fp-framing. Read counting and normalization were performed using fp-count and DESeq2, respectively. Translation efficiency (TE) and fold change values were calculated using the average values of replicates as follows (*i* indicates a gene):

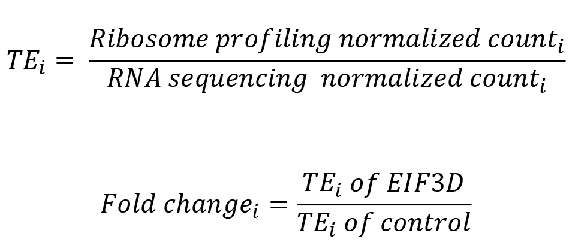

For identifying differentially translated transcripts, we employed DESeq2, utilizing a likelihood ratio test (model: Experiment + Target + Experiment:Target; reduced model: Experiment + Target). In pathway enrichment analysis, Enrichr was used with the WikiPathways_2021_Human database. To analyze upstream, downstream, and newly identified open reading frames (ORFs), we first obtained a bed file using ORF-RATER with ribosome profiling data of the parental iPSC line WTB6, treated with CHX and harringtonine^66,67^. Using this bed file, reads from control and sgEIF3D (Dox+ 3 days) samples were counted by fp-count. We then conducted a statistical analysis to identify significantly different transcripts between these conditions, employing DESeq2 with a Wald test.

## Statistics

The quantitative measurement results are presented as individual data points, depicted by colored dots, with means indicated by bars. In some instances, bar graphs with individual data points and error bars representing standard deviations are used. Statistical analyses included unpaired two-tailed t-tests to calculate p-values, assessing differences between two groups. Furthermore, one-way analysis of variance (ANOVA) was utilized for multiple comparisons. These analyses were performed using GraphPad Prism version 8.0.2 (GraphPad) and Excel (Microsoft). Statistical significance was determined by p-values or adjusted p-values less than 0.05, denoted by asterisks in the figures. The specific values are detailed in the figure legends.

### Key resources table

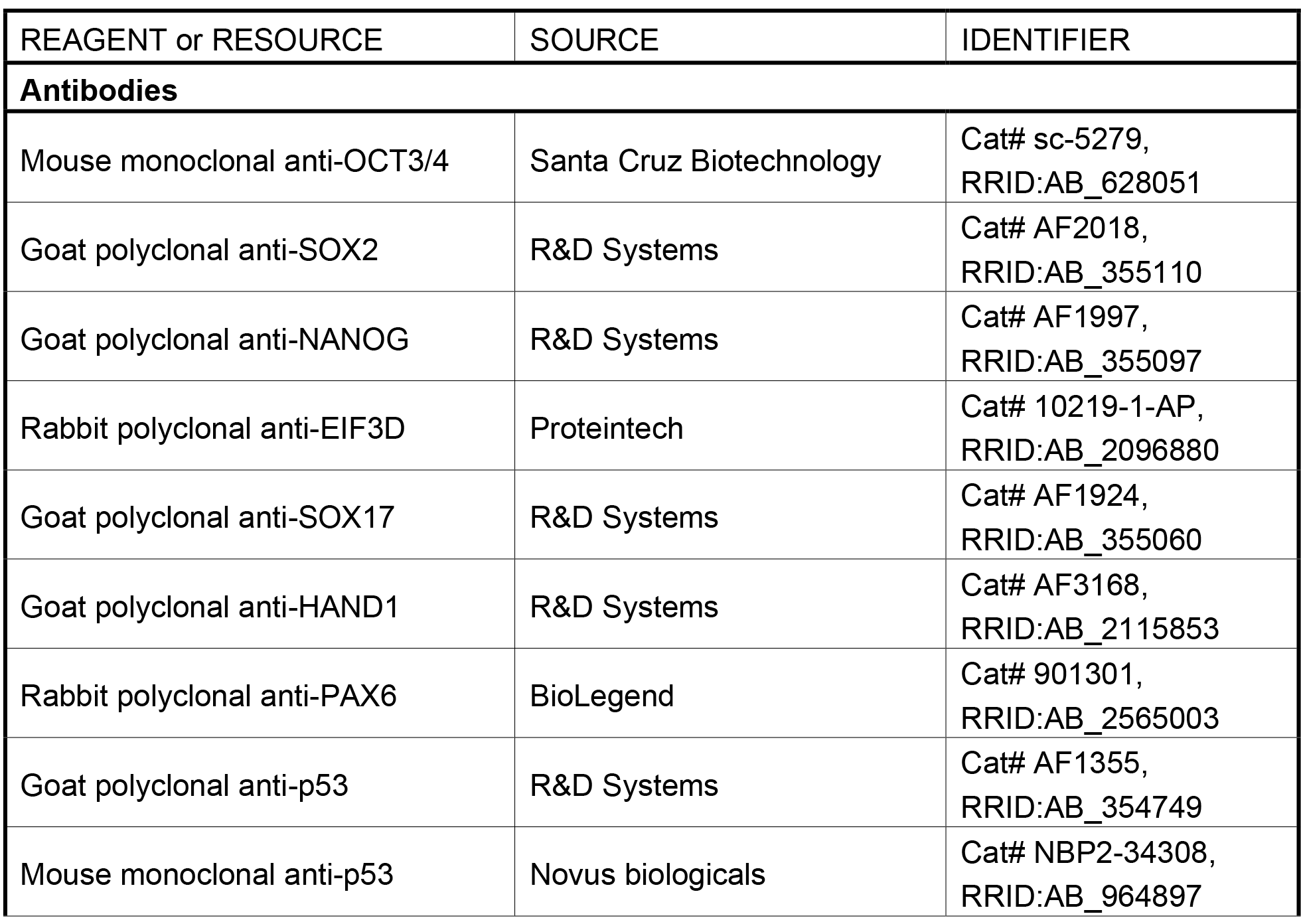
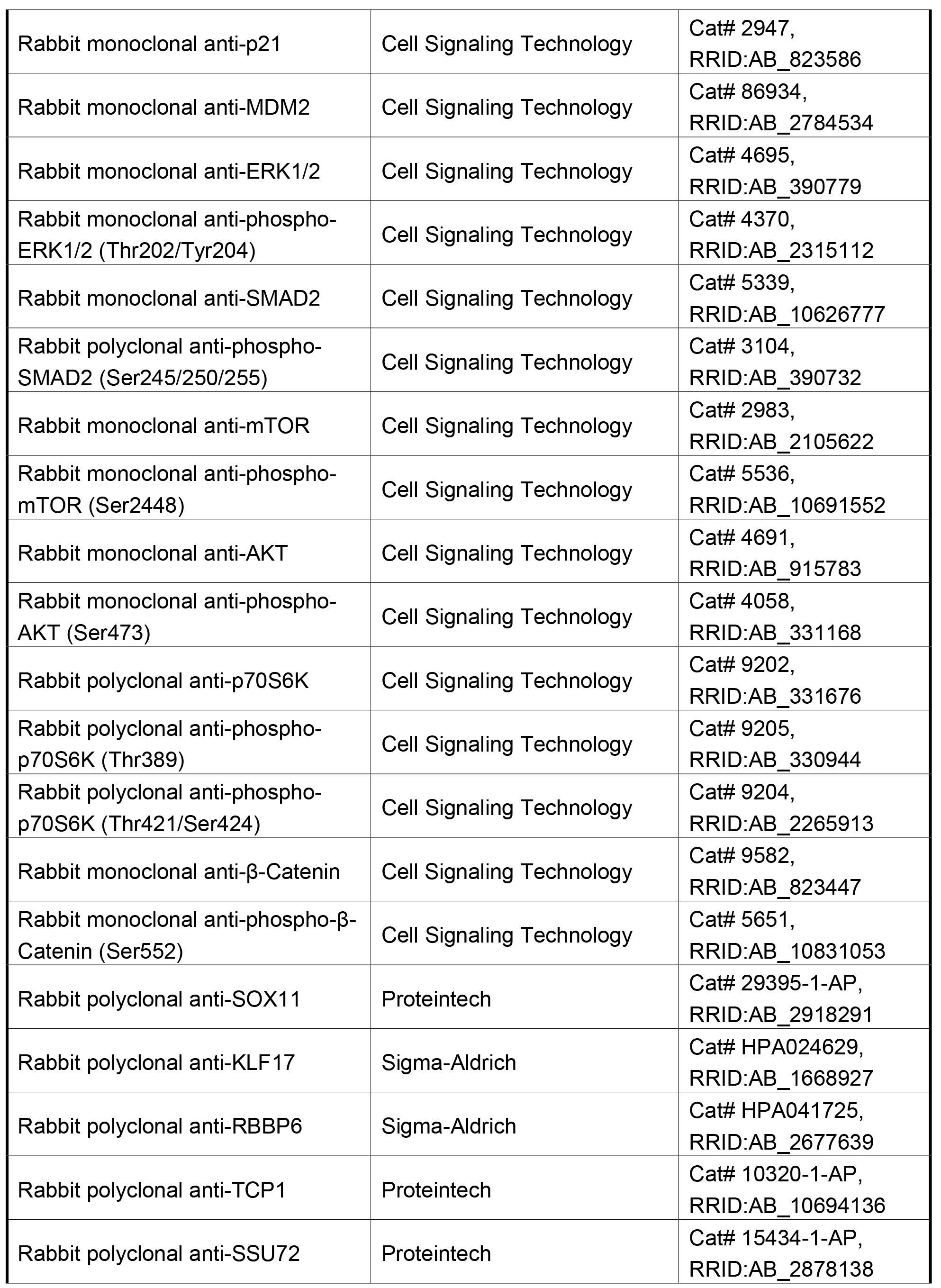
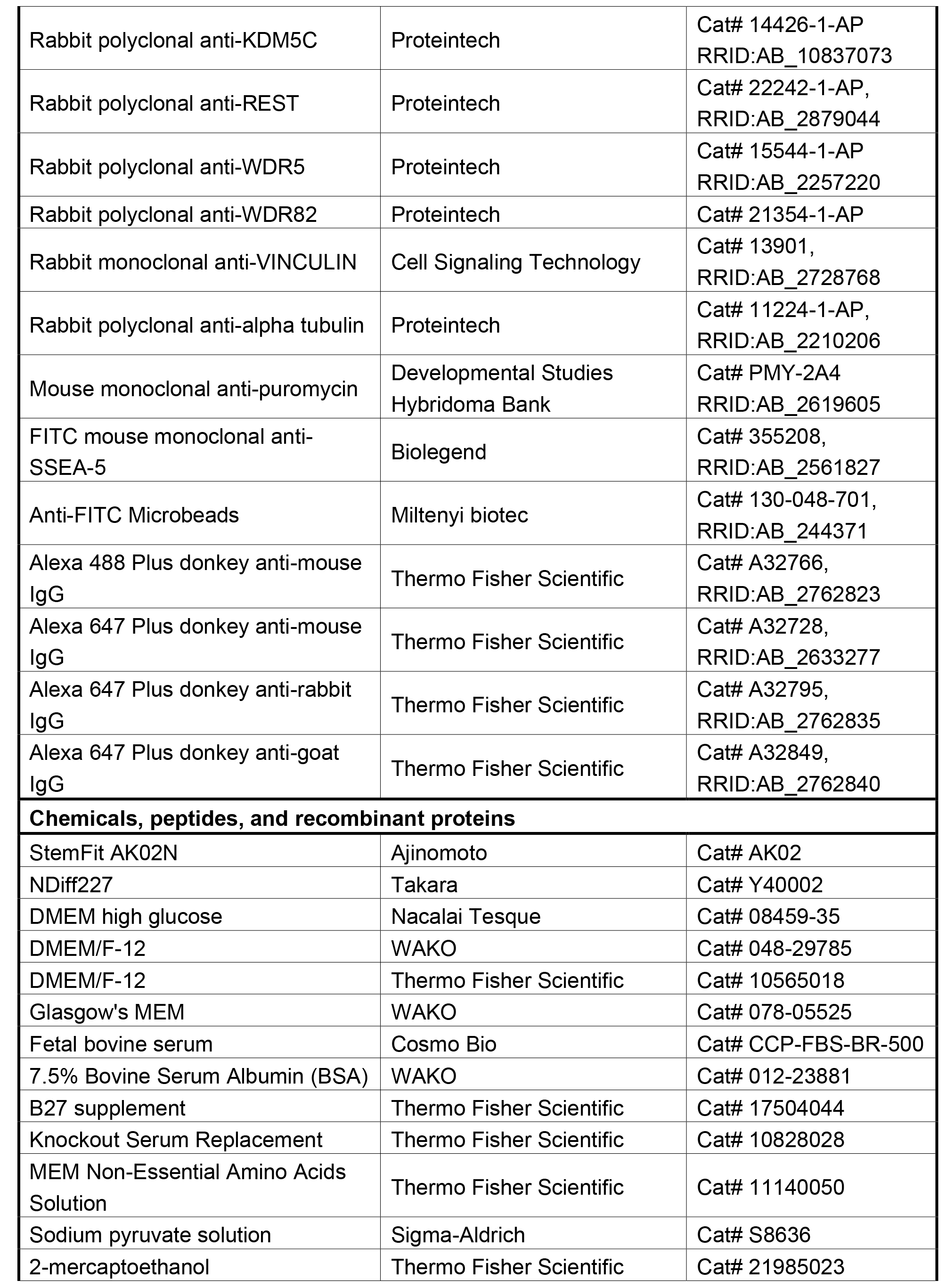
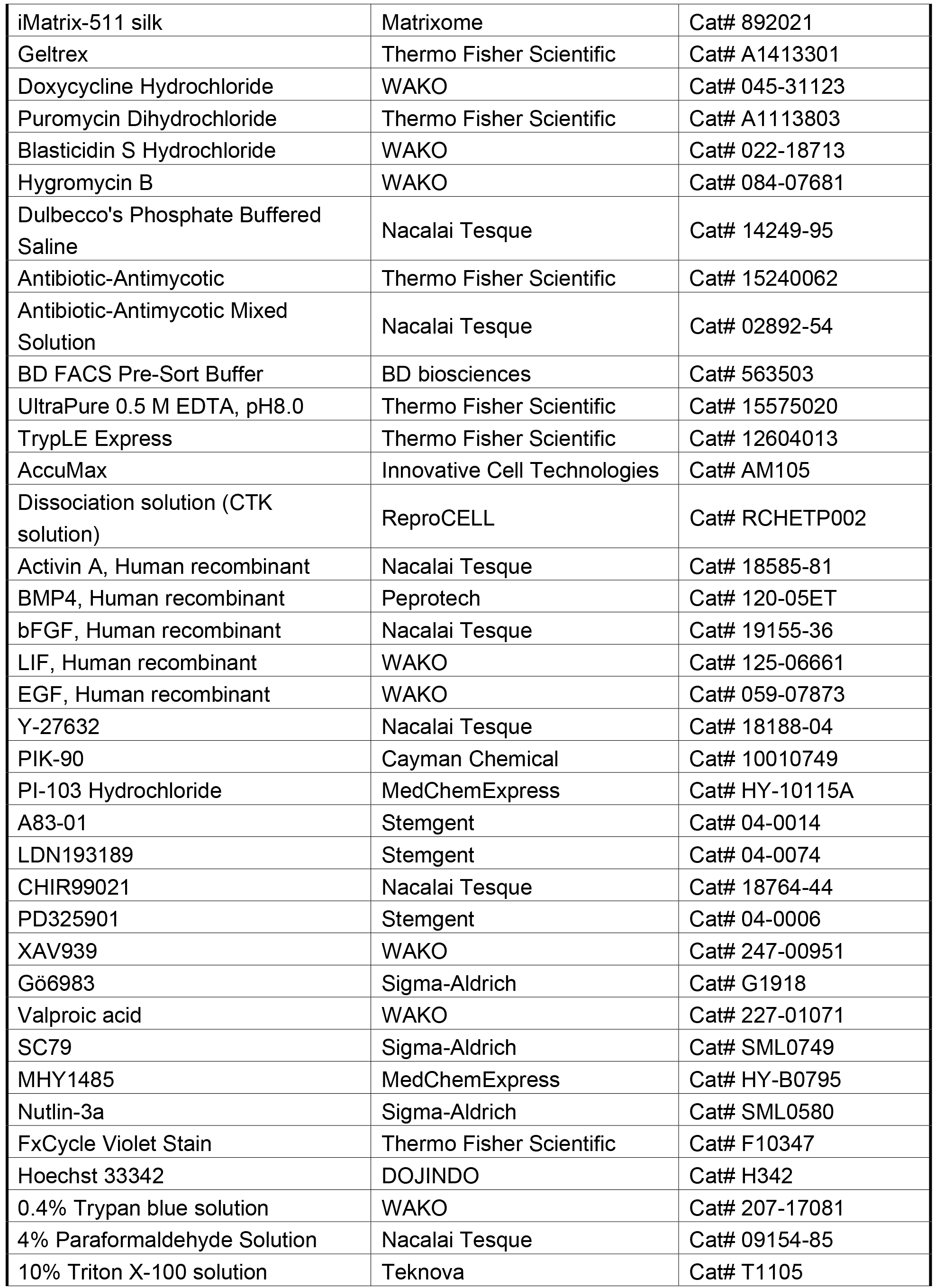
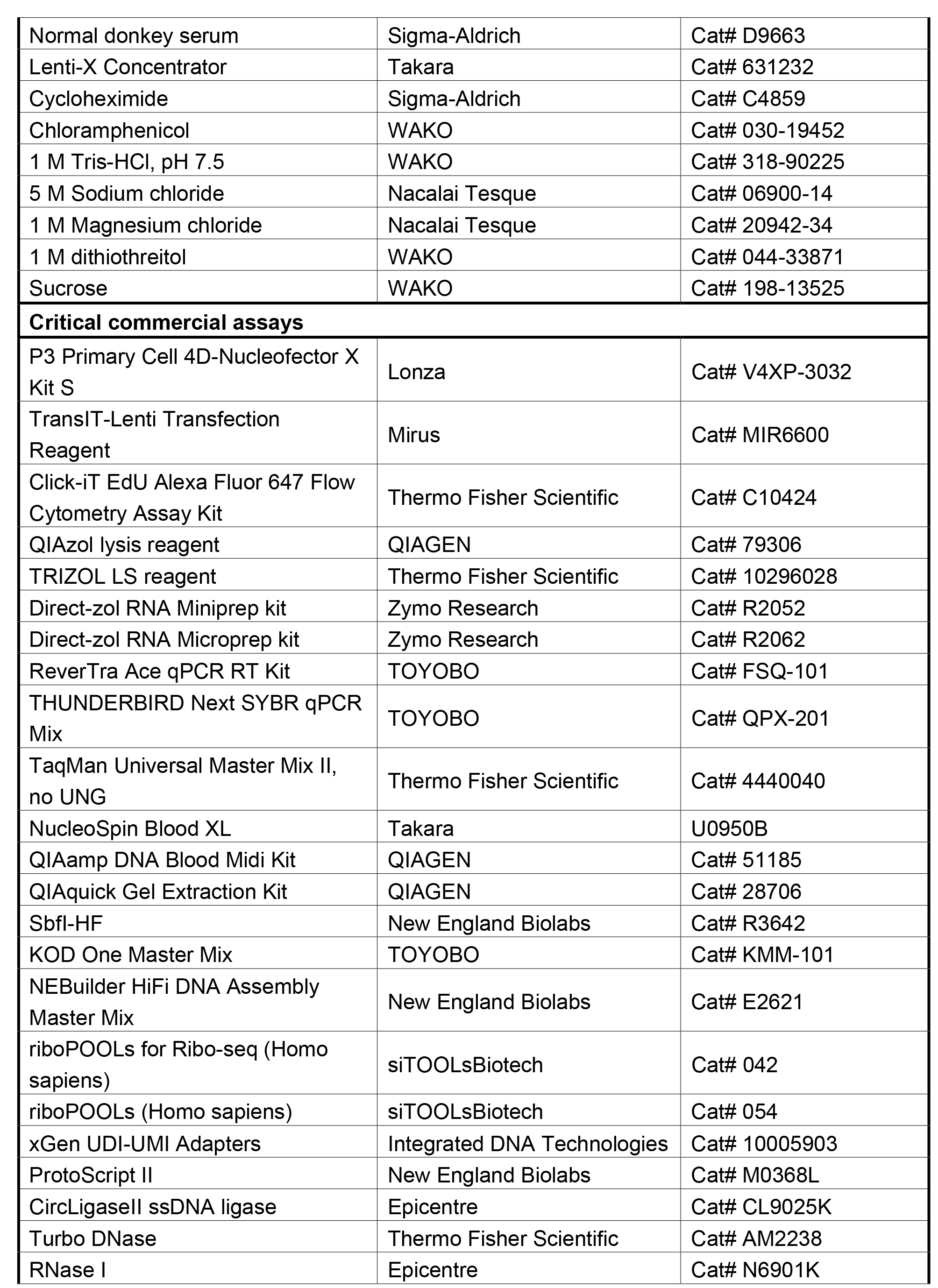
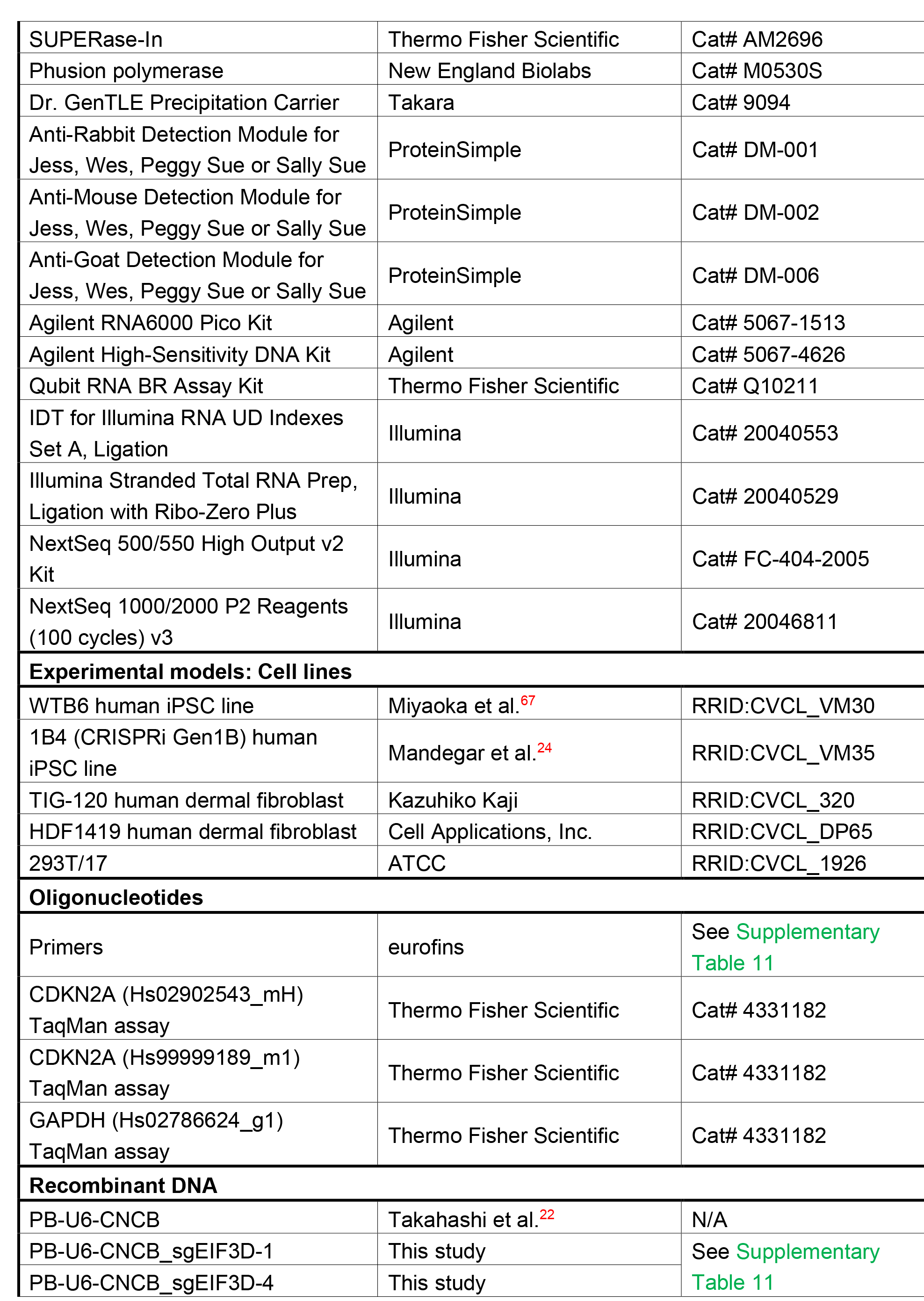
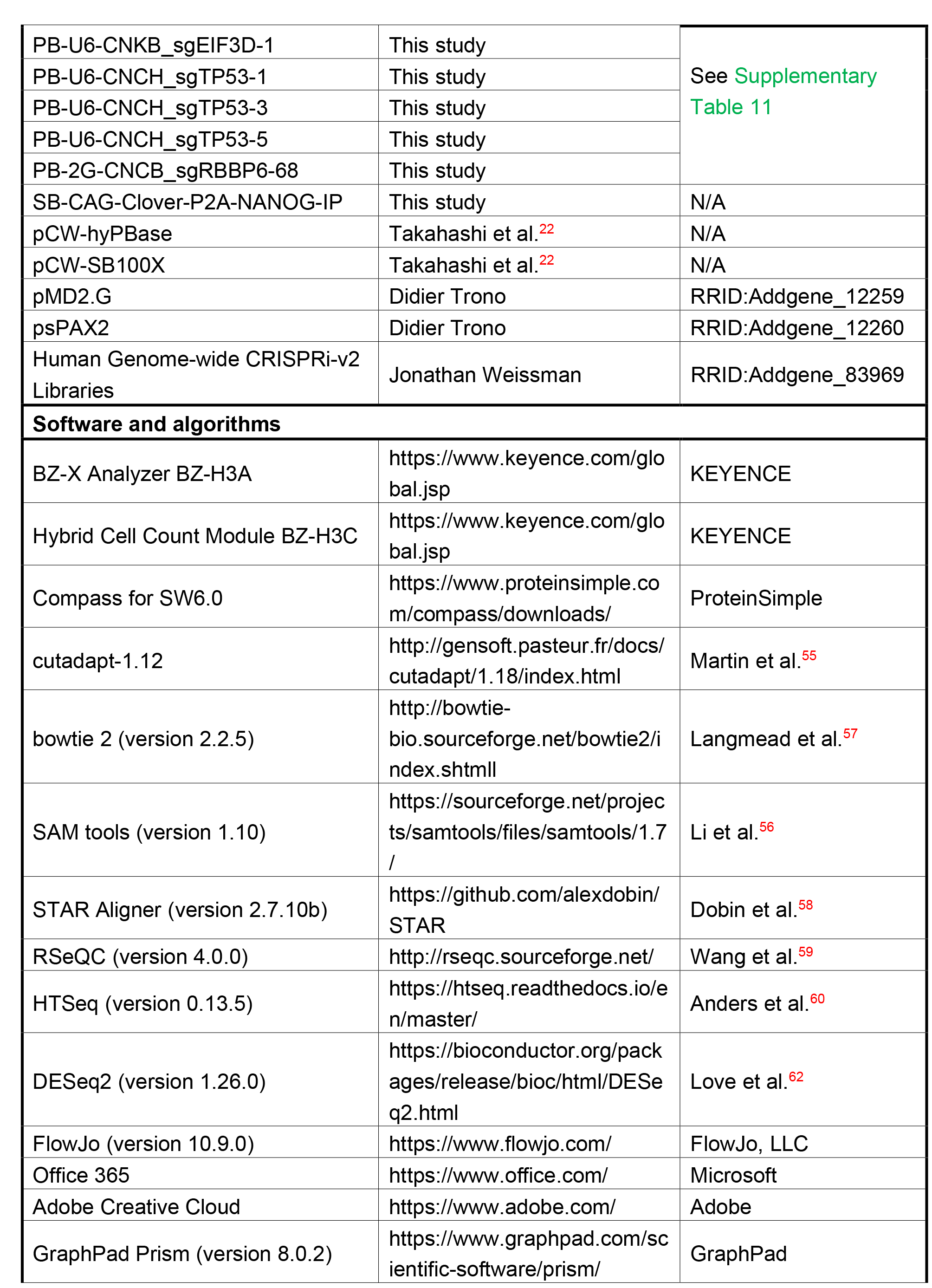
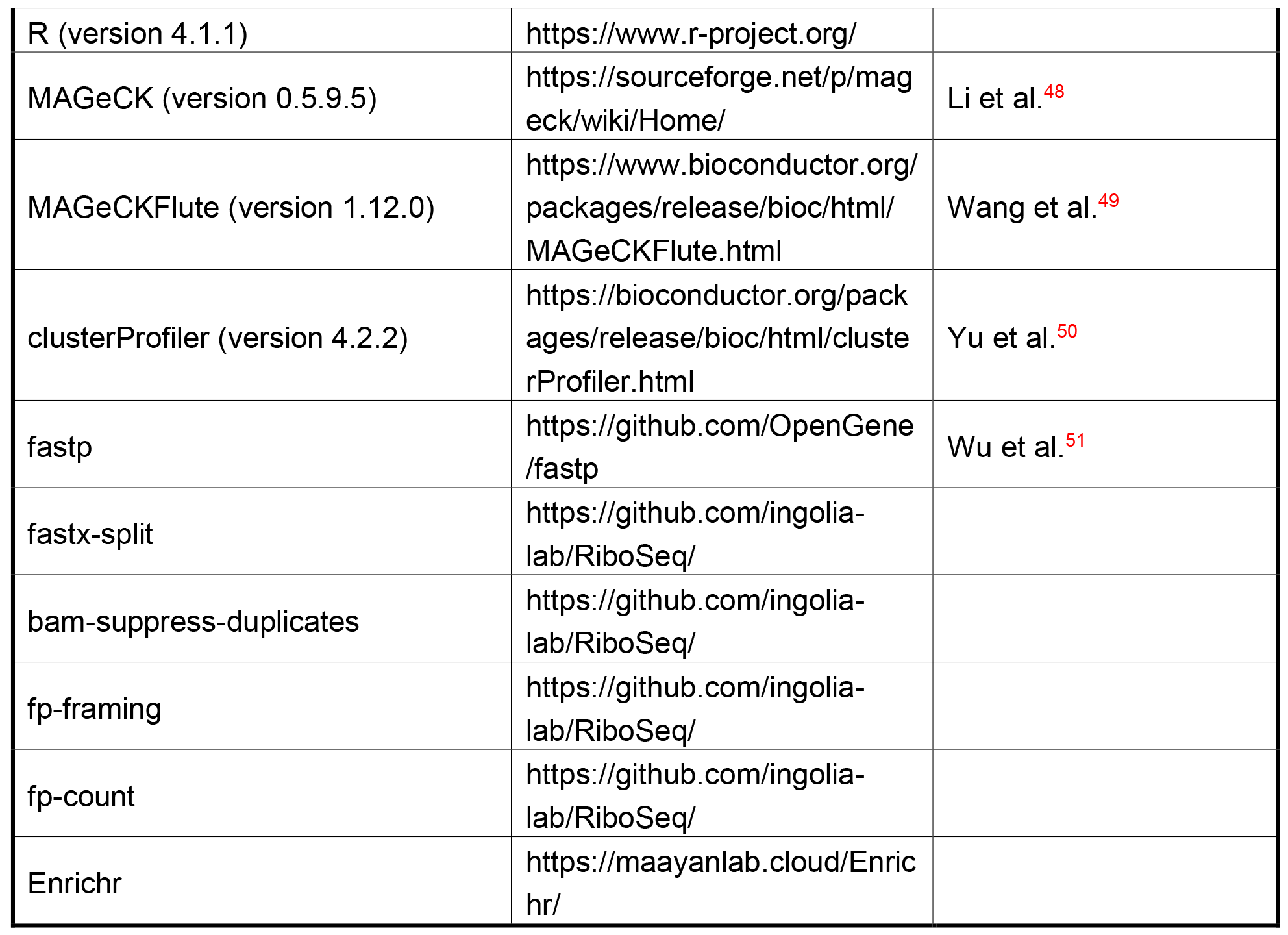

## Acknowledgments

We thank B. Conklin, M. Iwasaki, K. Kaji, M. Khurram, S. Perli, Y. Sato, K. Tomoda, D. Trono, J. Weismann, and S. Yamanaka for sharing materials and equipment, M. Hamao, K. Mitsunaga, A. Ogawa, and T. Wada for technical assistance and discussion, Y. Ishida and S. Takeshima for administrative support, and Editage for English language editing. This work was supported by Grants-in-Aid for Scientific Research from the Japanese Society for the Promotion of Science (JSPS) 20K20585 (KT), Grants-in-Aid for Scientific Research from JSPS 21H02155 (KT), Grants- in-Aid for Scientific Research from JSPS 22K14829 (CO), ACT-X grant from Japan Science and Technology Agency JPMJAX2222 (CO), Core Center for Regenerative Medicine and Cell and Gene Therapy from Japan Agency for Medical Research and Development (AMED) JP23bm1323001 (KT), Core Center for iPS Cell Research from AMED JP21bm0104001 (KT), a grant from the Takeda Science Foundation (KT), a grant from the Uehara Memorial Foundation (KT), a grant from the Mochida Memorial Foundation (KT), and the iPS Cell Research Fund from the Center for iPS Cell Research and Application, Kyoto University (CO, KT).

## Author Contributions

Conceptualization: C.O., K.T.; Methodology: C.O., M.S., Y.S., M.M., Y.T., S.I., K.T.; Investigation: C.O., M.N., K.T.; Visualization: C.O., K.T.; Funding acquisition: C.O., K.T.; Project administration: C.O., K.T.; Supervision: K.T.; Writing – original draft: C.O., K.T.; Writing – review & editing: all authors

---

**Supplementary Fig. 1:**
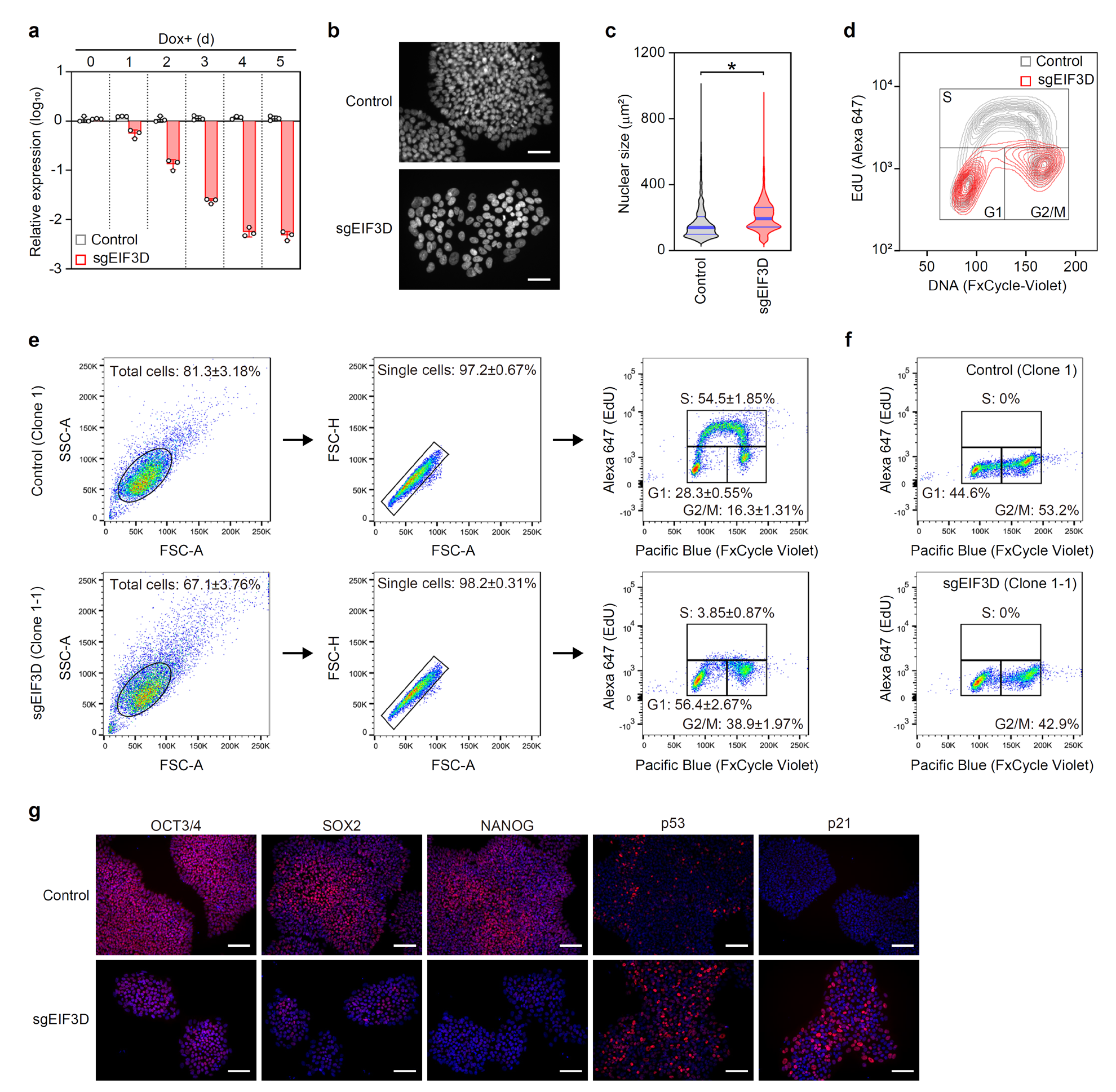
Supporting Data for EIF3D KD in Primed PSCs (Related to. Fig. 1**). a**, EIF3D expression on specified days post-KD induction, normalized to GAPDH and compared to control on day 0 (mean ± SD, n=3). **b**, Representative images of Hoechst 33342-stained control and sgEIF3D primed PSCs, 5 days post-KD induction. Scale bars: 100 μm. **c**, Nuclear size in control (n=5287) and sgEIF3D (n=5257) iPSCs, 5 days after Dox addition. p=1.18e-47, determined by unpaired t-test. **d**, Flow cytometry panels showing DNA content and EdU incorporation in control (grey) and sgEIF3D (red) primed PSCs, 5 days post-KD induction. **e**, Standard gating strategy used in flow cytometry analysis. **f**, Flow cytometry data of control (upper) and sgEIF3D (lower) primed PSCs, 5 days post-KD induction, without EdU incorporation. **g**, Immunocytochemistry images of control (upper) and sgEIF3D (lower) primed PSCs, 5 days post- KD induction, showing specified proteins (red). Nuclei visualized with Hoechst 33342 (blue). Scale bars: 100 μm.

**Supplementary Fig. 2:**
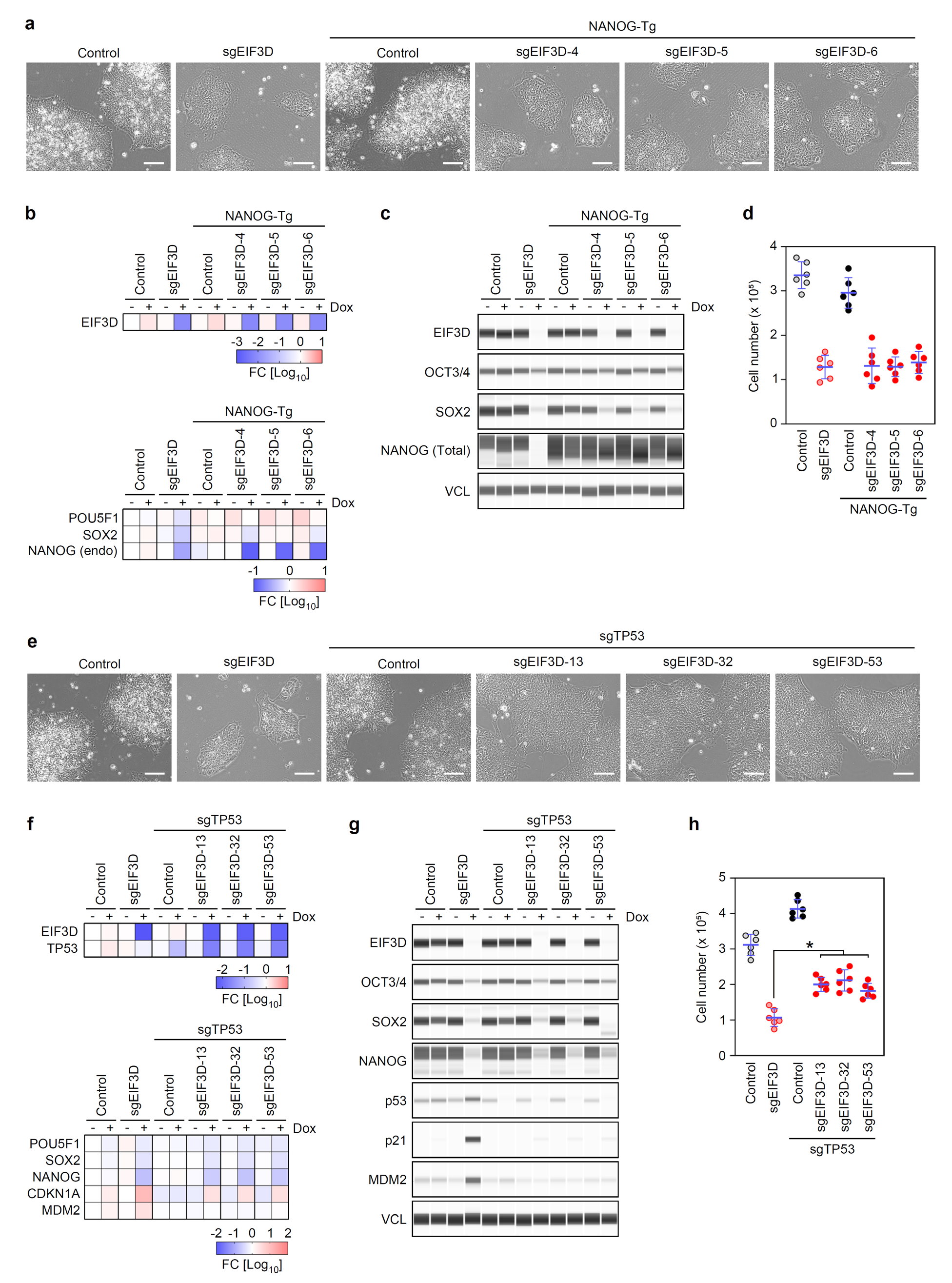
Impact of NANOG Overexpression and TP53 KD in EIF3D KD Primed PSCs (Related to Fig. 1). **a**, Representative images of control and sgEIF3D primed PSCs, with and without NANOG transgene (Tg), 5 days post-KD induction. Scale bars: 100 μm. **b**, Relative expression of EIF3D and pluripotency markers in cells from Supplementary Fig. 2a, normalized to GAPDH and compared to the control without Dox. n=3. **c**, Protein expression of EIF3D and pluripotency markers in cells depicted in Supplementary Fig. 2a. **d**, Cell counts from Supplementary Fig. 2a, 5 days post-KD induction (mean ± SD, n=6). **e**, Representative images of control and sgEIF3D primed PSCs, with and without sgTP53, 5 days post-KD induction. Scale bars: 100 μm. **f**, Relative expression of specified transcripts in cells from Supplementary Fig. 2e. Values normalized to GAPDH and compared to control without Dox. n=3. **g**, Protein expression in cells depicted in Supplementary Fig. 2e. **h**, Cell counts from Supplementary Fig. 2e, 5 days post-KD induction (mean ± SD, n=6). Control vs. sgTP53: p<0.0001; Control+sgTP53 vs. sgEIF3D+sgTP53: p<0.0002, determined by one-way ANOVA.

**Supplementary Fig. 3:**
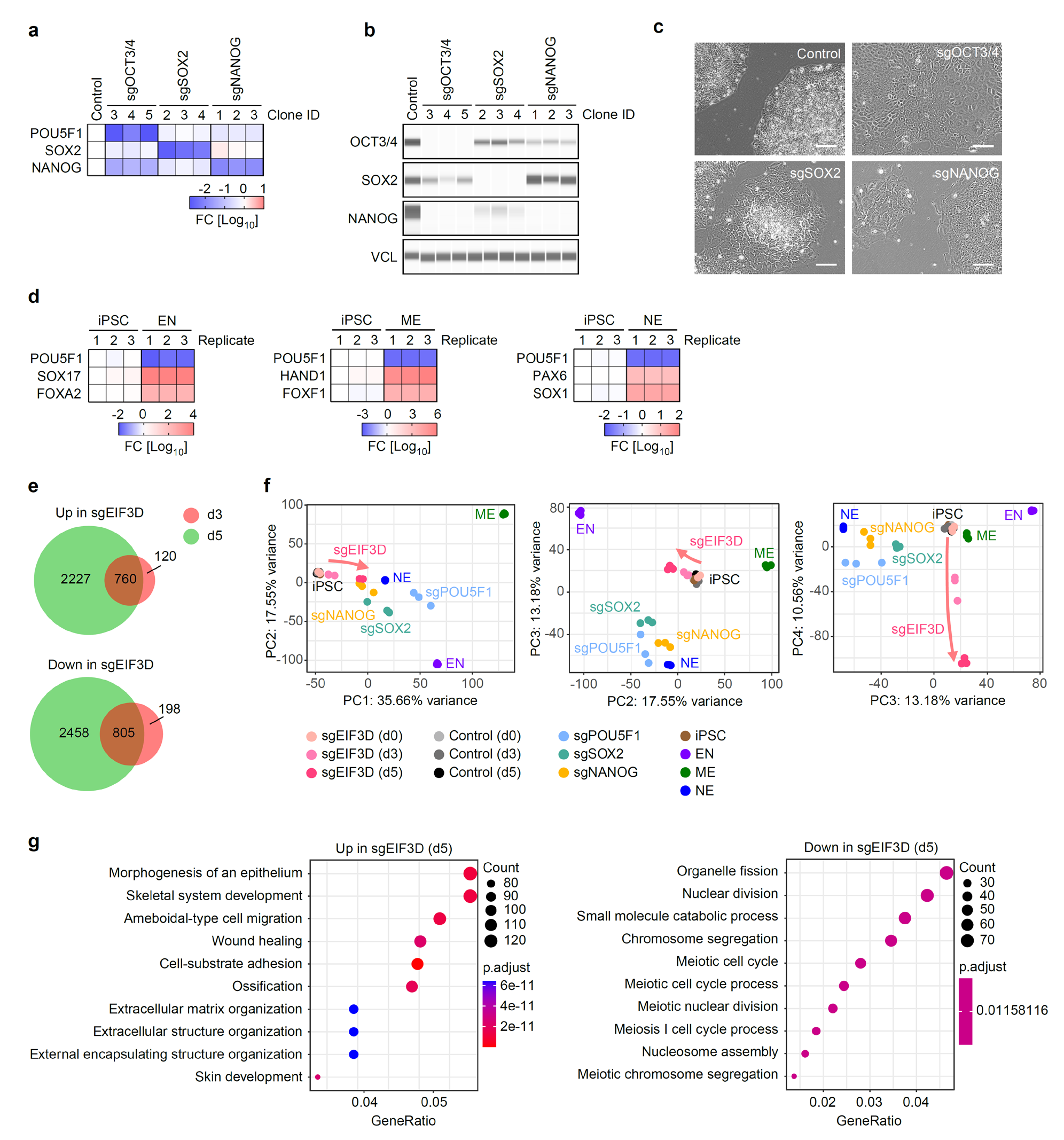
Transcriptome Changes Induced by EIF3D Knockdown in Primed PSCs (Related to Fig. 2). **a**, Relative expression of pluripotency markers in cells expressing sgPOU5F1, sgSOX2, and sgNANOG, 5 days post-Dox addition. Values are normalized to GAPDH and compared with control+Dox. n=3. **b**, Expression of pluripotency marker proteins in the cells depicted in Supplementary Fig. 3a. **c**, Representative images of the indicated cells, 5 days post-KD induction. Scale bars: 100 μm. **d**, Relative expression of POU5F1 as a pluripotency marker and various lineage markers in cells differentiated into specified lineages. Values are normalized to GAPDH and compared with undifferentiated 1B4 iPSCs (P35). n=2. **e**, Venn diagrams illustrating the overlap of differentially expressed genes (DEGs) on days 3 (top) and 5 (bottom) post-KD induction. **f**, Panels displaying the PCA results of RNA-seq from Fig. 2a, detailed on indicated principal component (PC) axes. **g**, Gene Ontology (GO) analyses for the DEGs.

